# Oxysterols drive inflammation via GPR183 during influenza virus and SARS-CoV-2 infection

**DOI:** 10.1101/2022.06.14.496214

**Authors:** Cheng Xiang Foo, Stacey Bartlett, Keng Yih Chew, Minh Dao Ngo, Helle Bielefeldt-Ohmann, Buddhika Jayakody Arachchige, Benjamin Matthews, Sarah Reed, Ran Wang, Matthew J. Sweet, Lucy Burr, Jane E. Sinclair, Rhys Parry, Alexander Khromykh, Kirsty R. Short, Mette Marie Rosenkilde, Katharina Ronacher

## Abstract

**Rationale:** Severe viral respiratory infections are often characterized by extensive myeloid cell infiltration and activation and persistent lung tissue injury. However, the immunological mechanisms driving excessive inflammation in the lung remain elusive.

**Objectives:** To identify the mechanisms that drive immune cell recruitment in the lung during viral respiratory infections and identify novel drug targets to reduce inflammation and disease severity.

**Methods:** Preclinical murine models of influenza virus and SARS-CoV-2 infection.

**Results:** Oxidized cholesterols and the oxysterol-sensing receptor GPR183 were identified as drivers of monocyte-macrophage infiltration to the lung during influenza virus (IAV) and SARS-CoV-2 infections. Both IAV and SARS-CoV-2 infections upregulated the enzymes cholesterol 25-hydroxylase (CH25H) and cytochrome P450 family 7 subfamily member B1 (CYP7B1) in the lung, resulting in local production of the oxidized cholesterols 25-hydroxycholesterol and 7α,25-dihydroxycholesterol (7α,25-OHC). Loss-of-function mutation of GPR183, or treatment with a GPR183 antagonist, reduced macrophage infiltration and inflammatory cytokine production in the lungs of IAV- or SARS-CoV-2-infected mice. The GPR183 antagonist also significantly attenuated the severity of SARS-CoV-2 infection by reducing weight loss and viral loads.

**Conclusion:** This study demonstrates that oxysterols drive inflammation in the lung and provides the first preclinical evidence for therapeutic benefit of targeting GPR183 during severe viral respiratory infections.

**Author Summary:** Viral infections trigger oxysterol production in the lung, attracting macrophages via GPR183. Blocking GPR183 reduced inflammation and disease severity in SARS-CoV-2 infection, making GPR183 a putative target for therapeutic intervention.

## Introduction

Severe viral respiratory infections including influenza and COVID-19 are associated with extensive myeloid cell recruitment to the lung, which can lead to a cytokine storm, severe tissue injury and the development of acute respiratory distress syndrome (ARDS) (1, 2). A shift in lung macrophage composition and function is associated with COVID-19 severity. A study of >600 hospitalised patients found that in severe cases resident alveolar macrophages were depleted and replaced by large numbers of inflammatory monocytes and monocyte-derived macrophages (3). Rapid monocyte infiltration of the lung during the acute phase of severe acute respiratory coronavirus 2 (SARS-CoV-2) infection is replicated in several animal models (4–6). On the other hand, monocyte recruitment is also an essential component of repair following lung injury (7). Therapeutic approaches are required that balance pro- inflammatory and pro-repair functions of recruited monocytes.

Oxidized cholesterols, so called oxysterols, have recently emerged as markers of inflammation in the lung. Oxysterols were increased in bronchoalveolar lavage fluid (BALF) from inflamed airways after allergen challenge and strongly correlated with infiltrating leukocytes (8). They were also increased in the sputum from patients with chronic obstructive pulmonary disease (COPD) correlating with disease severity (9, 10) and in the lungs of mice after lipopolysaccharide (LPS)-induced lung inflammation (9). However, the role of oxysterols in the lung during viral respiratory infections has not been investigated.

Oxysterols have a range of properties and receptors sharing a common role in inflammation (11, 12). One of these oxysterol producing pathways leads to the production of 7α,25-hydroxycholesterol (7α,25-OHC), via cholesterol 25-hydroxylase (CH25H) and cytochrome P450 family 7 subfamily B member 1 (CYP7B1) (12, 13) (**Figure 1A**). 7α,25-OHC is the endogenous high affinity agonist of the oxidized cholesterol-sensing G protein-coupled receptor GPR183 (also known as Epstein-Barr virus-induced gene 2; EBI2) (14, 15). GPR183 is expressed on cells of the innate and adaptive immune systems, including macrophages, dendritic cells, innate lymphoid cells, eosinophils and T and B lymphocytes (8, 16–18). With its oxysterol ligands GPR183 facilitates the chemotactic distribution of immune cells to secondary lymphoid organs (12, 14, 16, 17). *In vitro* GPR183 mediates migration of human and mouse macrophages towards a 7α,25-OHC gradient (19–21).

**Figure 1.**
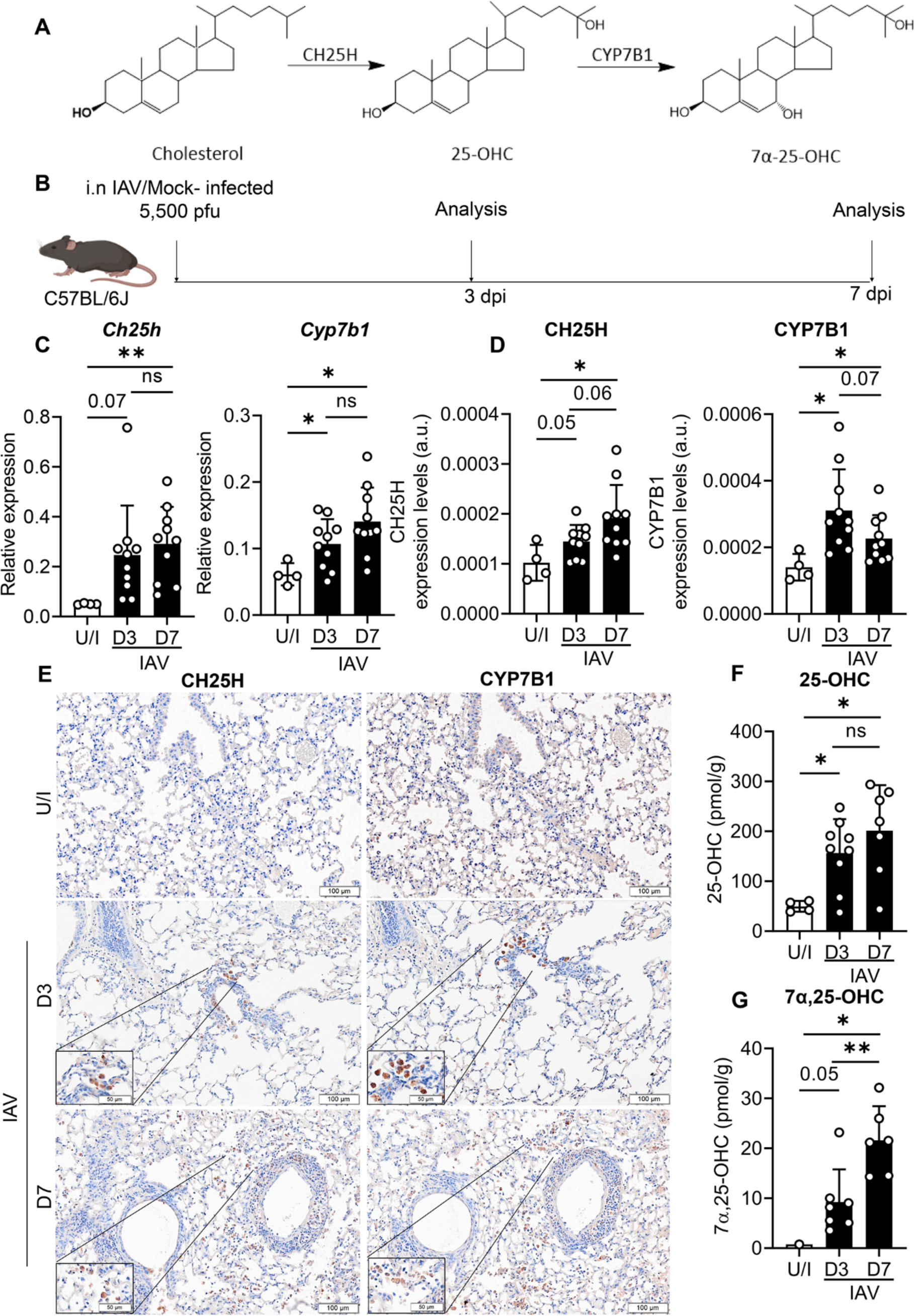
IAV infection leads to upregulation of CH25H and CYP7B1 expression in the lung and production of the oxysterols 25-OHC and 7α,25-OHC. **A**) The biosynthetic pathway of 25-OHC and 7α,25-OHC. **B**) Experimental design. C57BL/6J mice were infected intranasally with 5,500 PFU of A/Auckland/01/09 and mRNA expression of **C**) *Ch25h* and *Cyp7b1* were measured by qRT-PCR at 3 dpi and 7 dpi normalized to *Hprt*. **D**) Quantitative analysis of CH25H and CYP7B1 protein labelling by IHC. **E**) Representative IHC images of CH25H and CYP7B1 in lung sections of uninfected or IAV-infected mice. Concentrations of **F**) 25-OHC and **G**) 7α,25-OHC in the lungs at 3 dpi and 7 dpi expressed in pmol per gram lung tissue. Data are presented as mean ± SD of n=4 uninfected and n=6-10 infected mice per timepoint. Scale Bar = 100μm; dpi = days post-infection; U/I = mock infected; ns., not significant; *, *P* < 0.05; **, *P* <0.01 indicate significant differences.

In this study, we hypothesized that viral respiratory infections lead to the production of oxysterols in the lung and that these oxysterols contribute to excessive immune cell infiltration and inflammation. We show here that oxysterols drive GPR183-dependent monocyte infiltration in preclinical models of IAV and SARS-CoV-2 infection. Administration of a GPR183 antagonist significantly reduces inflammation, viral load and disease severity in mice infected with SARS-CoV-2. Accordingly, GPR183 is a putative host target for therapeutic intervention to mitigate disease severity in viral respiratory infections.

## Methodology

### Ethics and biosafety

All experiments were approved by the University of Queensland Animal Ethics Committee (MRI-UQ/596/18, AE000186) by the Institutional Biosafety Committee of the University of Queensland (IBC/465B/MRI/TRI/AIBN/2021).

### Viral Strains

Virus stocks of A/H1N1/Auckland/1/2009(H1N1) (Auckland/09) were prepared in embryonated chicken eggs. Viral titers were determined by plaque assays on Madin-Darby canine kidney (MDCK) cells as previously described (22). A mouse-adapted SARS-CoV-2 strain was obtained through serial passage of SARS-CoV-2 (B.1.351; hCoV-19/Australia/QLD1520/2020, GISAID accession EPI_ISL_968081, collected on 29 December 2020, kindly provided by Queensland Health Forensic and Scientific Services). Six x 10^4^ PFU of B1.351 was administrated intranasally to ketamine-anesthetized mice. Mice were monitored daily for weight loss and clinical signs of disease severity. Four days after inoculation, mice were euthanized, and bronchoalveolar lavage (BAL) was performed. The BALF was subsequently pooled and used to intranasally inoculate a new batch of mice. The process was repeated until a virulent phenotype of the virus was observed as determined by weight loss and clinical signs, which happened after four passages. To determine whether the mouse adapted SARS-CoV-2 acquired mutations sequencing of viral RNA was performed. Briefly, viral RNA was extracted from BALF using the Qiagen Mini kit and the quality confirmed suing the Agilent Bioanalyzer with 210 Expert software. Library preparations was performed using the Illumina Stranded Total RNA Ribo Zero Plus kit. Sequencing was performed using the NextSeq Midoutput kit, 125bp paired-end configuration with 19-25 million reads per sample. Sequencing analysis was executed using Galaxy software. Whole-genome ssequencing revealed a C to T mutation in position 10804 of the SARS-CoV-2 Beta genome resulting in the NSP5 mutation P252L. This mutation was rapidly selected from 3.4% in the initial virus stock to 8.8% in passage one. From passage two, this mutation reached consensus (60%) and underwent further fixation in passage three at 87% to final frequency of 92% in passage four. A mutation in NSP5 was detected in this mouse adapted SARS-CoV-2 strain (**Figure S1**). BALF of the mice from the fourth passage was subsequently pooled and used to inoculate Vero E6 cells for propagation, creating the viral stocks for our mouse-adapted strain. To verify the virulent phenotype of the mouse-adapted virus was retained after propagation in Vero E6 cells, the cell grown virus was used to inoculate a new batch of mice. The same viral stock was used to infect mice with 8x10^4^ PFU for the experiments described.

### Bioinformatic analysis of mouse-adapted SARS-CoV-2 sequence data

Base-called fastq files were mapped to the QLD1520 SARS-CoV-2 isolate (GISAID accession EPI_ISL_968081) using Bowtie2 (v2.4.2) (Langmead and Salzberg 2012) under default alignment conditions. Sub consensus variants of alignment files were identified using iVar (v1.2.2) (Grubaugh et al. 2019) with a minimum quality score threshold of 20 and depth of 5000. Coverage of mapped alignment files was determined using samtools (v1.3) depth. Frequencies and coverage of variant positions were manually validated using Integrative Genomics Viewer (Version: 2.7.0) (Thorvaldsdottir, Robinson, and Mesirov 2013). Variant frequencies and alignment depth was 4isualized using GraphPad Prism (v9.3.1). Raw fastq data generated in this study have been deposited in the Sequence Read Archive hosted by the National Center for Biotechnology Information with accession number PRJNA849351.

### Plaque assays

IAV plaque assays were carried out on confluent monolayers of MDCK cells as previously described (22). SARS-CoV-2 plaque assays were carried out on Vero E6 cells as described previously (23).

### Mouse models

*Gpr183*tm1Lex were obtained from Lexicon Pharmaceuticals (The Woodlands, USA), back-crossed to a C57BL/6J background and bred in-house at the Biological Resources Facility at the Translational Research Institute, Australia. Eight to 10-week-old C57BL/6J and *Gpr183*tm1Lex (C57BL/6J background; *Gpr183*^-/-^) mice were anesthetized with isoflurane (4% isoflurane, 0.4 L/min oxygen flow rate) before being inoculated intranasally with 5,500 PFU of A/Auckland/01/09 (H1N1). Mice were monitored for weight loss. For SARS-CoV-2 infection, C57BL/6J and *Gpr183^-/-^* mice were sanesthetized with ketamine/Xylanzine (80mg/kg/5mg/kg) before being inoculated intranasally with 8x10^4^ PFU of mouse-adapted SARS-CoV-2 and monitored for weight loss. Lungs were collected at specified timepoints for subsequent downstream analysis. The GPR183 antagonist NIBR189 was administrated from 1 dpi. IAV infected mice were sacrificed at 3 dpi and 7 dpi for examination. SARS-CoV-2 infected mice were sacrificed at 2 dpi and 5 dpi. Lungs homogenised in DMEM for use in plaque assays and ELISAs. For RNA processing, lungs were collected in TRIzol (Invitrogen). For oxysterol extraction, lungs were collected in methanol. For histological analysis the lungs were fixed in 10% neutral buffered formalin.

### RNA isolation and RT-qPCR

Total RNA was isolated using ISOLATE II RNA Mini Kit (Bioline Reagents Ltd., London, UK) as previously described (24). The list of primers (Sigma Aldrich) is provided in **Table S1**. The relative expression (RE) of each gene using the 2^-ΔCt^ method, normalizing to the reference gene (Hypoxanthine-guanine phosphoribosyltransferase; HPRT).

### Oxysterol extraction from lung tissues

The oxysterol extraction and quantification method was adapted from Ngo et al. (24). Lung lobes from IAV and SARS-CoV-2-infected mice were homogenized in methanol. Oxysterols were extracted using a 1:1 dichloromethane:methanol solution containing 50 µg/mL BHT in a 30°C ultrasonic bath. Tubes were flushed with nitrogen to displace oxygen, sealed with a polytetrafluoroethylene (PTFE)-lined screw cap, and incubated at 30°C in the ultrasonic bath for 10 mins. Following centrifugation (3,500 rpm, 5 min, 25°C), the supernatant from each sample was decanted into a new tube. For liquid-liquid extraction, Dulbecco’s phosphate-buffered saline (DPBS) was added to the supernatant, agitated and centrifuged at 3,500 rpm for 5 mins at 25°C. The organic layer was recovered and evaporated under nitrogen using a 27-port drying manifold (Pierce; Fisher Scientific, Fair Lawn, NJ). Oxysterols were isolated by solid-phase extraction (SPE) using 200 mg, 3 mL aminopropyl SPE columns (Biotage; Charlotte, NC). The samples were dissolved in 1 ml of hexane and transferred to the SPE column, followed by a rinse with 1 ml of hexane to elute nonpolar compounds. Oxysterols were eluted from the column with 4.5 ml of a 23:1 mixture of chloroform: methanol and dried under nitrogen. Samples were resuspended in 50µl of warm (37°C) 90% methanol with 0.1% DMSO, and placed in an ultrasonic bath for 5 min at 30°C. A standard curve was extracted for 25-OHC (Sigma-Aldrich, H1015) and 7α,25-OHC (SML0541, Sigma-Aldrich) using the above method. Dichloromethane, butylated hydroxytoluene (BHT) and hexane were purchased from Sigma-Aldrich.

### Mass spectrometric quantitation of 25-OHC and 7α,25-OHC

Samples were analysed on an AB Sciex QTRAP® 5500 (ABSCIEX, Redwood City, CA) mass spectrometer coupled to a Shimadzu Nexera2 UHPLC. A Kinetex Pentafluorophenyl (PFP) column (100 × 2.1mm, 1.7µM, 100^0^A, Phenomenex) was used for the separation of 25-OHC and 7α,25-OHC from other oxysterols. Mobile phase used for separation were, A - 0.1% formic acid with water and B - 100% acetonitrile with 0.1% formic acid. Five µL of sample were loaded at 0.5 mL/min and separated using linear gradient with increasing percentage of acetonitrile. Samples were washed for 1.3 min after loading with 30% mobile phase B followed by linear gradient of 30% - 70% over 9 min and 70% to 99% over 1 min. The column was washed with 99% mobile phase B for 2 min followed by equilibration with 30% B 2 min before next injection. Column oven and auto-sampler were operated at 50°C and 15°C, respectively. Elution of analytes from the column was monitored in positive ion mode (ESI) with multiple reaction monitoring on ABSciex QTRAP® mass spectrometer equipped with Turbo spray ion source, which was operated at temp 550°C, ion spray voltage of 5500 V, curtain gas (CUR) of 30 psi, ion source gas1 (GS1) of 65 psi and ion source gas 2 (GS2) of 50 psi. Quadrupole 1 and 3 were operated at unit mass resolution at all time during the experiment. MRM pairs 385.3 > 367.3, 385 >133, 385.3 > 147.1 were monitored for 25-OHC and for 7α,25-OHC following MRM pairs were used 383.2 > 365.3, 383.2 > 147.3, 383.2 >159.0. Deuterated 25-OHC (11099, Sapphire Bioscience, Redfern, Australia) and 7α,25-OHC (700078P, Merck) were used as internal standards. Following MRM transitions were recoded for internal standards 391.1 > 373.2, 391.1 >133.1, 391.1>123.1 (25-OHC) and 407.2 > 389.0 (7α,25-OHC). De-clustering potential (DP), collision energy (CE), entrance (EP) and collision cell exit potential (CXP) were optimised for each MRM pair to maximise the sensitivity. Data was processed using AbSciex MultiQuant™ software (Version 3.0.3). Oxysterol concentrations were subsequently normalized to the lung weights. High-performance liquid chromatography (HPLC) grade methanol, acetonitrile and chloroform were purchased from Merck.

### Cytokine quantification using ELISA

Cytokines in lung homogenates were measured with DuoSet ELISA (IFNβ (DY8234-05), IFNγ (DY485), IFNλ (DY1789B), IL-6 (DY406), TNFα (DY410), IL-1β (DY401), IL-10 (DY417) and CCL2 (DY479), R&D systems) according to the manufacturer’s protocol.

### Flow cytometry

Lung lobes of IAV-infected mice digested in digestion buffer (Librase; Roche) and passed through 40-µm nylon mesh to obtain single cell suspensions. Red blood cells lysis performed using BD Pharm Lyse (BD Biosciences, San Jose, CA). Cells were labelled with: Zombie Green Fixable Viability kit (423111, Biolegend), PerCP-CD45 (30-F11), Brilliant Ultraviolet 395-CD3e (145-2C11, BD Biosciences), Brilliant Violet (BV) 786-CD4 (L3T4, BD Biosciences), PE/Cyanine7-CD11b (M1/70), BV510-CD11c (N418), APC/Cyanine7-F4/80 (BM8), BV605-Ly6G (1A8, BD Bioscience), PE-B220 (RA3-6B2), BV421-I-A/I-E (M5/114.15.2), APC-Siglec-F (CD170, S17007L, BD bioscience) before flow cytometric analysis on the BD LSRFortessa X20. Post-acquisition analysis was performed using FlowJo software (TreeStar).

### Immunohistochemistry

Heat-induced epitope retrieval was performed using citrate buffer (pH 6, 95°C, 30 mins) (BP327-1; Thermo Fisher Scientific). Sections were blocked for endogenous peroxidase activity using 3% hydrogen peroxide (HL001-2.5L-P, Chem Supply, Adelaide, South Australia), washed with tris-buffered saline (TBS; Bio-Rad) containing 0.05% polysorbate 20 (Tween-20; Sigma Aldrich; TBST) and blocked using background sniper (BS966, Biocare Medical, Concord, CA) for 30 mins. Immunohistochemistry (IHC) was performed on deparaffinized and rehydrated lung sections. Immunolabeling was performed with rabbit antibodies against SARS-CoV-2 nucleocapsid protein antibody (1 hour at 25°C, 1:5000) (40143-R040 Sino Biological), IBA1 (2 hours at 25°C 1:1000) (019-19741; NovaChem), CH25H (4°C overnight 1:600) (BS-6480R, Bioss Antibodies), CYP7B1 (4°C overnight 1:1000) (BS-5052R, Bioss Antibodies) and isotype control (rabbit IgG 31235, Thermo Fisher Scientific) diluted in Da Vinci Green Diluent (PD900, Biocare Medical) followed by incubation with horseradish peroxidase (HRP)-conjugated goat anti-rabbit Ig antibody (1:200) (ab6721, Abcam). Isotype controls are shown in (**Figure S2**). Sections were washed with TBST before applying chromogen detection, using diaminobenzidine (ab64238, DAB substrate kit Abcam,) as per the manufacturer’s instructions. Counterstaining was performed with Mayer’s hematoxylin (Sigma-Aldrich) before dehydrating the sections in a series of increasing ethanol concentrations (70% to 100% ethanol). Sections were clarified with xylene, and mounted using a xylene-based mounting medium (15-184-40, SHURMount Mounting Media, Fisher scientific). Slides were scanned in an Olympus SLIDEVIEW VS200 using a 20x objective. DAB-positive areas were quantified using ImageJ (https://imagej.nih.gov/ij/).

### Statistical analysis

Data were analysed on GraphPad Prism software. Data were also assessed for normality using Shapiro-Wilk test. Spearman rank correlation was used to analyse correlations. For two group comparisons, parametric Student’s two-tailed t test was used for normally distributed data while nonparametric Mann-Whitney U test was used for skewed data that deviate from normality.

## Results

### IAV infection increases CH25H and CYP7B1 expression and oxysterol production in the lung

To investigate whether IAV infection induces the production of oxidized cholesterols, we infected mice with IAV (**Figure 1B**) and determined the mRNA expression of oxysterol producing enzymes in the lung. *Ch25h* and *Cyp7b1* mRNA was increased in the lungs of IAV-infected mice compared to uninfected animals (**Figure 1C**). Similarly, CH25H and CYP7B1 protein expression was also increased, as demonstrated by immunohistochemical labelling of lung sections with antibodies detecting CH25H and CYP7B1 protein (**Figure 1D,E**). The induction of oxysterol producing enzymes was associated with increased concentrations of the oxysterols 7α,25-OHC and 25-OHC in IAV-infected lungs at both 3 days post infection (dpi) and 7 dpi (**Figure 1F, G**). In uninfected lungs, 7α,25-OHC was undetectable in most samples tested. Consistent with the increase in oxysterols, *Gpr183* mRNA was increased at 3 dpi and 7 dpi (**Figure S3A**), suggesting increased recruitment of GPR183-expressing immune cells to the lung upon infection. *Gpr183* expression was positively correlated with *Ch25h* and *Cyp7b1* (**Figure S3B, C**).

### *Gpr183*^-/-^ mice have reduced macrophage infiltration into the lungs upon IAV infection

To investigate whether oxysterol-mediated immune cell recruitment is dependent on oxysterol-sensing GPR183, we performed experiments in mice genetically deficient in *Gpr183* (*Gpr183^-/-^*). *Gpr183^-/-^* mice are viable and exhibit normal gross phenotype (25). However, upon infection with IAV, *Gpr183^-/-^* mice had lower IBA1^+^ macrophage numbers in the lung at 3 dpi and 7 dpi compared to infected C57BL/6J controls (**Figure 2A**). *Gpr183* expression was positively correlated with mRNA expression of the pro-inflammatory cytokines *Il6, Tnf* and *Ccl2* in C57BL/6J mice (**Figure S4**) and reduced macrophage infiltration in *Gpr183^-/-^* mice was associated with reduced *Il6* and *Tnf,* but not *Ccl2* at 7 dpi (**Figure S5**). Body weights and viral titers through the course of IAV infection were comparable across the two genotypes (**Figure S6**). These results demonstrate that GPR183 is required for macrophage infiltration into the lung upon IAV infection and that lower macrophage numbers are associated with reduced expression of pro-inflammatory cytokines.

**Figure 2.**
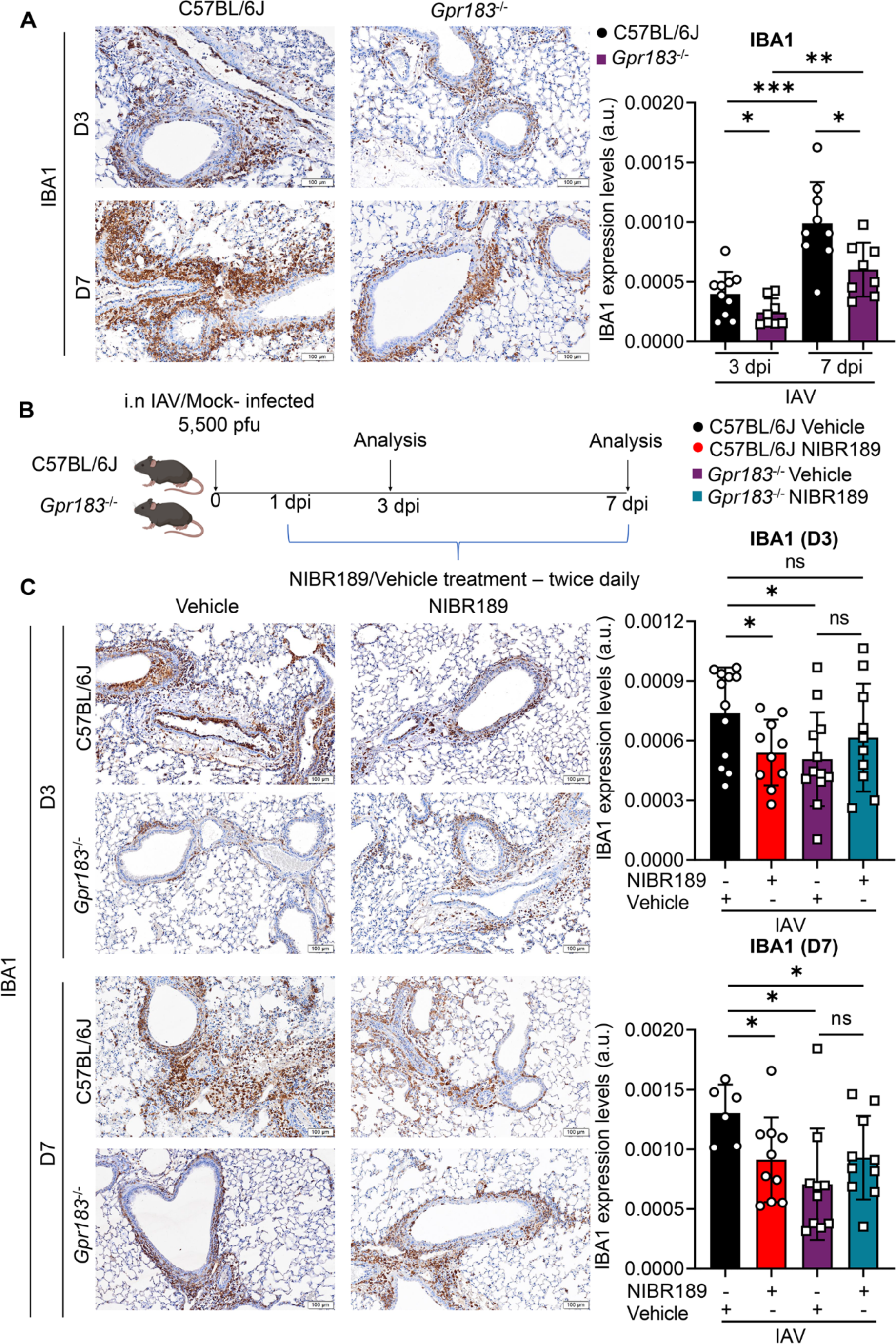
Deletion of the *Gpr183* gene or administration of a GPR183 antagonist reduces macrophage infiltration in IAV-infected lungs. C57BL/6J and *Gpr183^-/-^* mice were infected intranasally with 5,500 PFU of A/Auckland/01/09. **A**) Representative IHC images of IBA1 in lung sections of IAV-infected C57BL/6J and *Gpr183^-/-^*mice. Quantitative analysis of IBA1 staining. **B**) Experimental design; C57BL/6J mice and *Gpr183^-/-^* mice were infected intranasally with 5,500 PFU of A/Auckland/01/09. Mice were subsequently treated orally with 7.6 mg/kg NIBR189 or vehicle control twice daily from 1 dpi until the end of the experiment. **C**) Representative IHC images of IBA1 in lung sections of C57BL/6J and *Gpr183^-/-^* mice with the respective treatment groups at 3dpi and 7dpi. Quantitative analysis of IBA1 staining. Data are presented as mean ± SD of n = 6-12 infected mice per genotype and timepoint. dpi = days post-infection; Scale Bar = 100μm; U/I = mock infected ns = not significant; *, *P* < 0.05; **, *P* < 0.01; ***, *P* < 0.001 indicate significant differences

### GPR183 inhibition reduces macrophage infiltration

To investigate whether GPR183 is a putative therapeutic target to reduce inflammation, the synthetic GPR183 antagonist NIBR189 (14, 21) was administered to C57BL/6J mice twice daily starting from 24 h post-infection until the end of the experiment (**Figure 2B**). Like *Gpr183^-/-^* mice, C57BL/6J animals treated with NIBR189 had significantly reduced macrophage infiltration into the lung both at 3 and 7 dpi as determined by IHC (**Figure 2C**).

In addition, flow cytometry analysis was performed on lung single cell suspensions from C57BL/6J and *Gpr183^-/-^*mice treated with NIBR189 and vehicle, respectively, using a previously published gating strategy (26) (**Figure S7**). NIBR189-treated C57BL/6J mice and *Gpr183^-/-^* mice had lower percentages of macrophages (F480^high^/CD11b^+^/Ly6G^-^/SigF^-^) (**Figure 3A, B**) compared to vehicle-treated C57BL/6J animals after IAV infection. NIBR189 treatment did not change the percentages of other immune cell subsets in the lung, including neutrophils (B220^-^/CD3^-^/Ly6G^+^/CD11b^+^) (**Figure 3A, C**), CD4+ T cells, CD8+ T cells, B cells, DCs, and alveolar macrophages (**Figure S8**). Body weights and lung viral loads were not affected by genotype or treatment (**Figure S9**).

**Figure 3.**
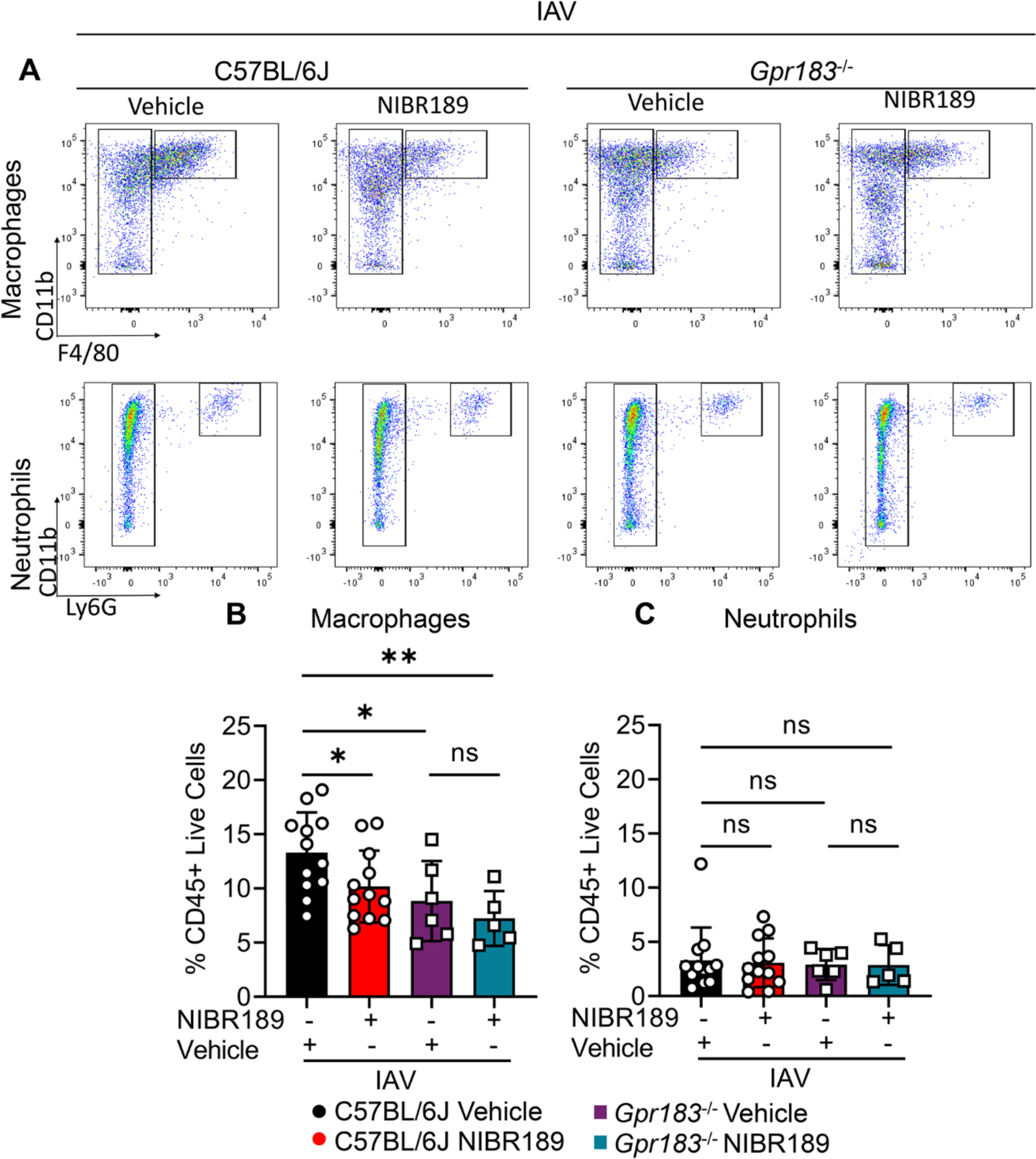
The GPR183 antagonist NIBR189 reduces macrophage infiltration and inflammatory cytokine production. C57BL/6J and *Gpr183^-/-^* mice were infected intranasally with 5,500 PFU of A/Auckland/01/09. Mice were subsequently treated orally with 7.6 mg/kg NIBR189 or vehicle control twice daily from 1 dpi until the end of the experiment. **A**) Frequency of infiltrating macrophages (F480^high^/CD11b^+^/Ly6G^-^/SigF^-^) and neutrophils (B220^-^ CD3^-^Ly6G^+^) was determined by flow cytometry relative to total viable CD45^+^ immune cells 3 dpi. Graphs depicting the frequency of **B**) macrophages and **C**) neutrophils. Data are presented as mean ± SD of n=5-12 infected mice per genotype and timepoint. dpi = days post-infection; U/I = mock infected; ns = not significant; *, *P* < 0.05; **, P<0.01 indicate significant differences.

Taken together our results demonstrate that the GPR183 antagonist NIBR189 significantly reduced the infiltration of macrophages to the lung without affecting the recruitment of other immune cell subsets to the site of infection.

### GPR183 inhibition reduces IAV-induced pro-inflammatory cytokine concentrations

We next determined if the reduced macrophage infiltration mediated by the GPR183 antagonist NIBR189 results in reduced inflammatory cytokine production in the lung. At 3 dpi, no significant differences in cytokine production were observed between treatment groups (**Figure S10**). However, IAV-Infected C57BL/6J mice treated with NIBR189 had significantly lower concentrations of IL-6, TNF and IFNβ (**Figure 4A-D**) at 7 dpi. This was again comparable to the phenotype of IAV-infected *Gpr183^-/-^* mice, with NIBR189 treatment having no additional effect in mice deficient in GPR183. In addition, no significant differences were observed in IFNλ across the two timepoints (**Figure 4D** **and Figure S10**) demonstrating that the GPR183 antagonist treatment does not negatively impact the production of type III IFNs which are important for viral control in the lung (27). No differences between treatment groups were observed at either timepoint for protein concentrations of IL-1β, CCL2 or IFNγ between treatment groups (**Figure S10 and S11**). Thus, GPR183 can be inhibited pharmacologically to reduce proinflammatory cytokines upon severe IAV infection.

**Figure 4.**
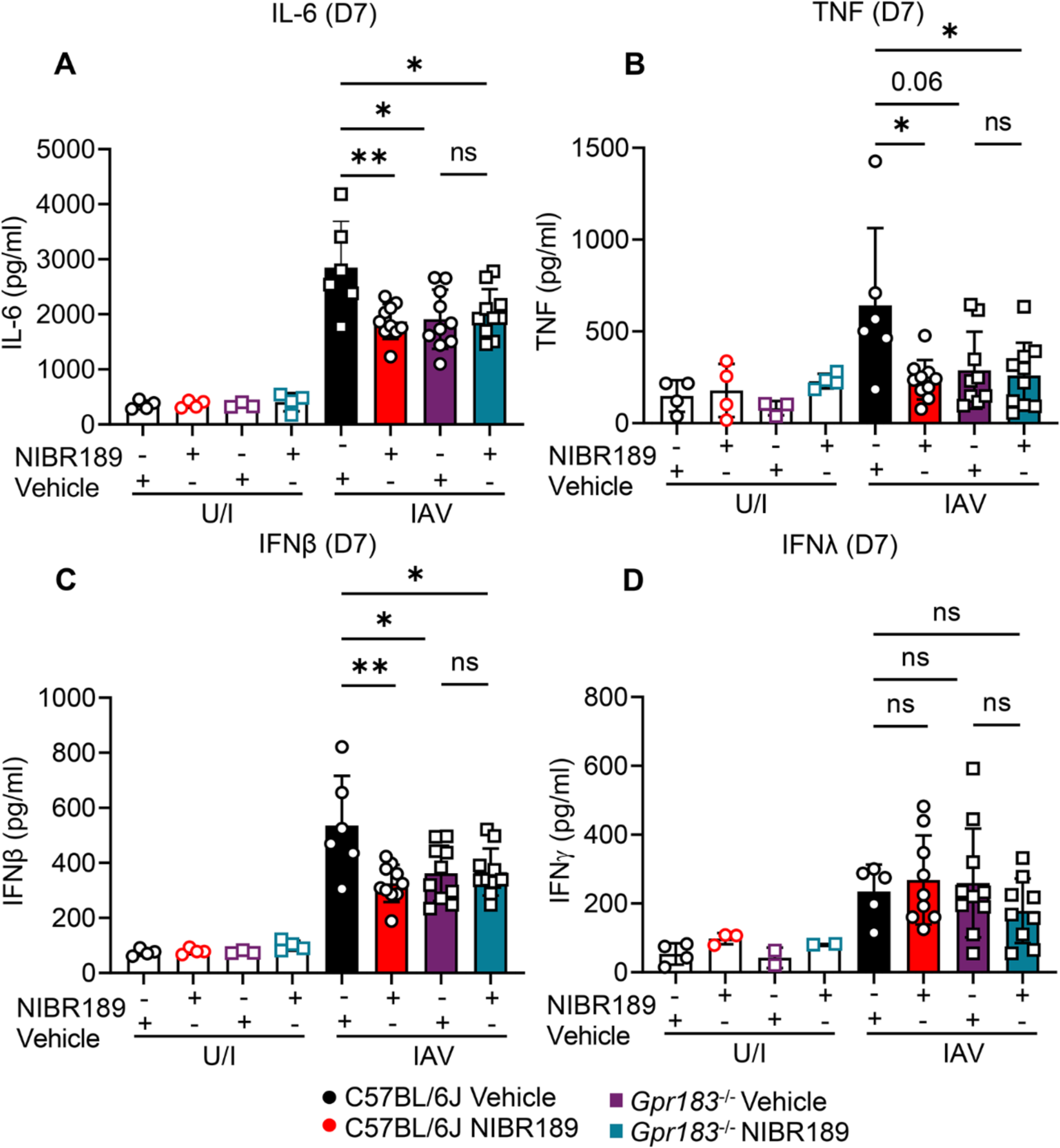
The GPR183 antagonist NIBR189 reduces inflammatory cytokine production. C57BL/6J and *Gpr183^-/-^* mice were infected intranasally with 5,500 PFU of A/Auckland/01/09. Mice were subsequently treated orally with 7.6 mg/kg NIBR189 or vehicle control twice daily from 1 dpi until the end of the experiment. Cytokine measurements of **A**) IL-6, **B**) TNF, **C**) IFNβ and **D**) IFNλ at 7 dpi measured by ELISA. Data are presented as mean ± SD of n=5-12 infected mice per genotype and timepoint. dpi = days post-infection; U/I = mock infected; ns = not significant; *, *P* < 0.05; **, P<0.01 indicate significant differences.

### GPR183 inhibition reduces SARS-CoV-2 infection severity

Excessive macrophage infiltration and activation is a hallmark of severe COVID-19 (3, 28). To evaluate whether the benefits of inhibiting GPR183 extend to SARS-CoV-2 infection, we established a mouse-adapted SARS-CoV-2 strain by passaging the Beta variant of SARS-CoV-2 (B.1.351) four times in C57BL/6J mice. This resulted in a viral stock that contained a mutation in NSP5 and caused clinical signs in infected mice as indicated by body weight loss (**Figure S1**). Consistent with the IAV infection results, mRNA expression of *Ch25h* and *Cyp7b1* was significantly upregulated in the lungs of SARS-CoV-2 infected mice compared to uninfected mice (**Figure 5A**). This was confirmed also at the protein level by IHC (**Figure 5B, C**). Further, 25-OHC and 7α,25-OHC concentrations in lung homogenates were significantly increased at 2 dpi, returning to uninfected levels by 5 dpi by which time the animals began to recover from the infection (**Figure 5D**). NIBR189 or vehicle was administered to C57BL/6J or *Gpr183^-/-^* mice twice daily from 24 h post-SARS-CoV-2 infection until the end of the experiment (**Figure 6A**). NIBR189-treated C57BL/6J mice lost significantly less weight and recovered faster compared to infected C57BL/6J mice receiving vehicle (**Figure 6B** **and S12**). Similarly, *Gpr183^-/-^* had less severe SARS-CoV-2 infection. Collectively, these data demonstrate that oxysterols are produced in the lung upon SARS-CoV-2 infection and inhibition of GPR183 significantly reduced the severity of SARS-CoV-2 infection.

**Figure 5.**
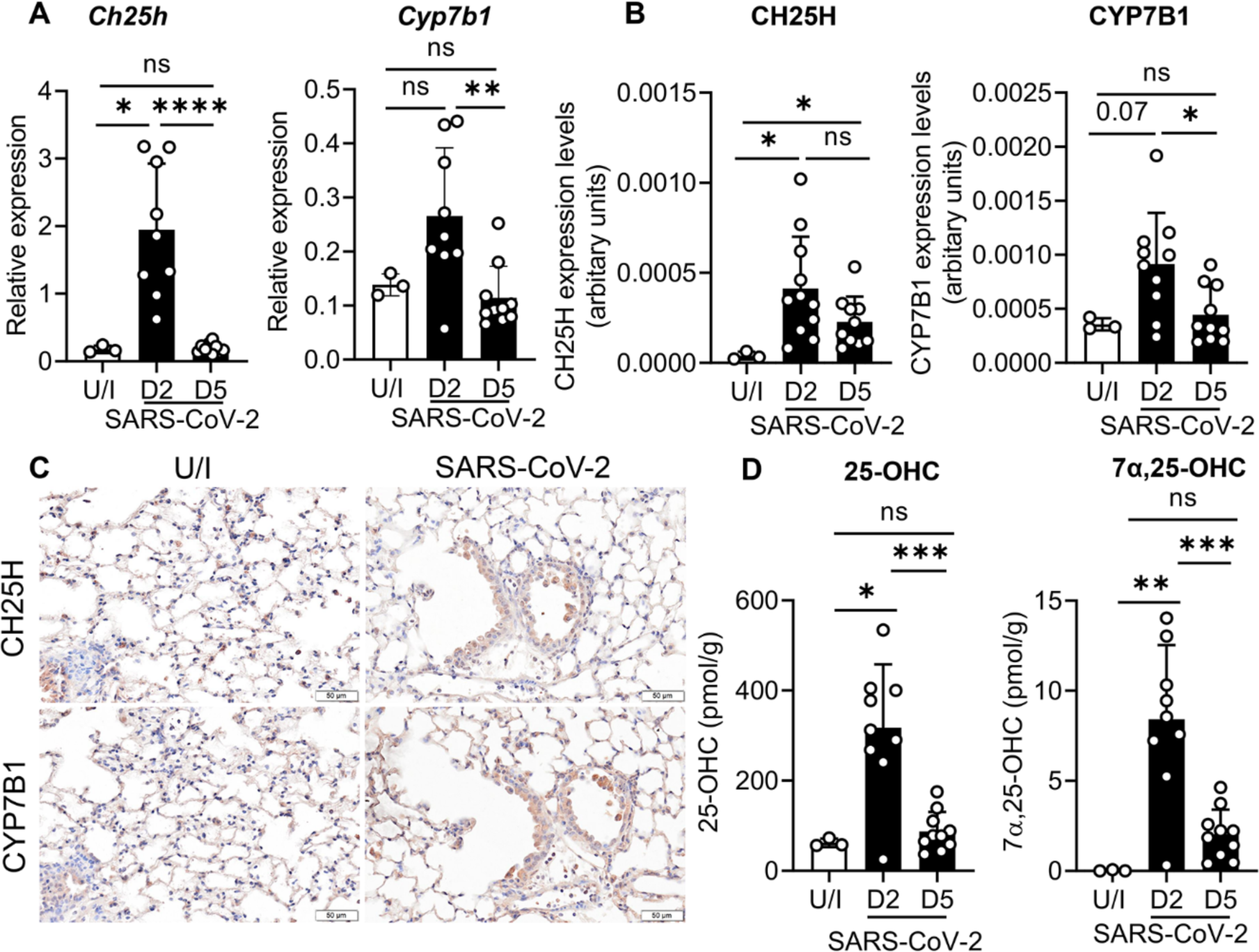
SARS-CoV-2 infection leads to upregulation of CH25H and CYP7B1 expression in the lung and production of the oxysterols 25-OHC and 7α,25-OHC. C57BL/6J mice were infected intranasally with approximately 8x10^4^ PFU of mouse-adapted SARS-CoV-2. mRNA expression of **A**) *Ch25h* and *Cyp7b1* was measured by qRT-PCR at 2 dpi and 5 dpi normalized to *Hprt*. **B**) Quantitative analysis of CH25H and CYP7B1 protein by IHC labelling and **C**) representative IHC images of CH25H and CYP7B1 in lung sections in uninfected, 2 dpi and 5 dpi. **D**) Concentrations of 25-OHC and 7α,25-OHC in the lungs at 2 dpi and 5 dpi expressed in pmol per gram lung tissue. Data are presented mean ± SD of n=3 uninfected mice and n= 9-10 infected mice per timepoint. Scale Bar = 50μm; U/I = mock infected; dpi = days post-infection; ns = not significant; *, *P* < 0.05; **, *P* < 0.01; ***, *P* < 0.001; ****, *P* < 0.0001 indicate significant differences.

**Figure 6.**
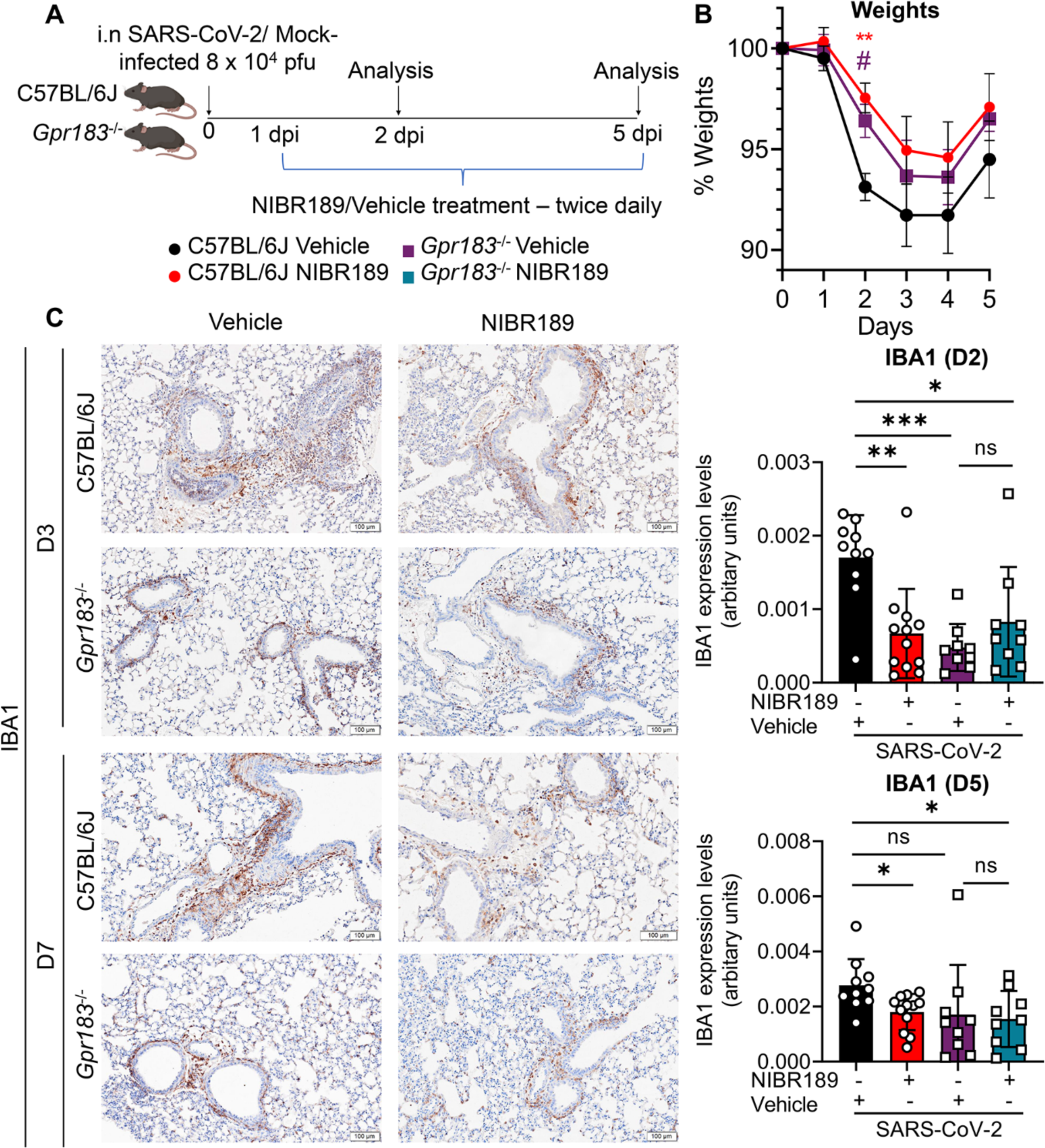
GPR183 inhibition resulted in less SARS-CoV-2 infection-induced weight loss and in reduced macrophage infiltration. C57BL/6J and *Gpr183^-/-^*mice were infected intranasally with approximately 8x10^4^ PFU of mouse-adapted SARS-CoV-2. Mice were subsequently treated orally with 7.6 mg/kg NIBR189 or vehicle control twice daily from 1 dpi until the end of the experiment. **A**) Experimental design. **B**) Weights of mice displayed as percentage of the weight at time of inoculation. **C**) Representative IHC images of IBA1 in lung of C57BL/6J and *Gpr183^-/-^* mice with the respective treatment groups at 2 dpi and 5 dpi (left). Scale Bar = 100μm. Quantitative analysis of IBA1 (right). Data are presented mean ± SD of n=9-12 infected mice per genotype and timepoint. Scale Bar = 100μm; U/I uninfected; dpi = days post-infection; ns = not significant; *, *P* < 0.05; **, *P* < 0.01; ***, *P* < 0.001 indicate significant differences.

### GPR183 inhibition reduces macrophage infiltration and inflammatory cytokine expression in the lung of SARS-CoV-2 infected mice

Next, we investigated whether the inhibition of GPR183 also decreases macrophage infiltration and inflammatory cytokines in the lung. SARS-CoV-2-infected C57BL/6J mice treated with NIBR189 had significantly reduced macrophage infiltration into the lung at 2 dpi and 5 dpi (**Figure 6C**). NIBR189 treatment was also associated with reduced *Tnf, Il10* and *Ifng* mRNA expression at 2 dpi (**Figure 7A-C**), as well as reduced *Tnf*, *Il1b and Il6* expression at 5 dpi (**Figure 7D-F**). Early interferon responses were not affected by NIBR189 treatment with comparable *Ifnb* and *Ifnl* expression at 2 dpi in C57BL/6J mice that received NIBR189 treatment versus vehicle (**Figure 8A, B**). Late interferon responses (5 dpi) were significantly lower in NIBR-treated animals compared to controls (**Figure 8C, D**). No differences between treatment groups were observed for mRNAs encoding *Ccl2, Il1b*, or *Il6* at 2 dpi as well as those encoding *Ccl2*, *Il10* and *Ifng* at 5 dpi (**Figure S13**). These results demonstrate that reduced macrophage infiltration in NIBR-treated mice was associated with reduced pro-inflammatory cytokine expression in the lung, while the early antiviral IFN responses remained unchanged. The mechanism(s) by which oxysterols attract macrophages to the lung to produce pro-inflammatory cytokines are therefore conserved across viral infections.

**Figure 7.**
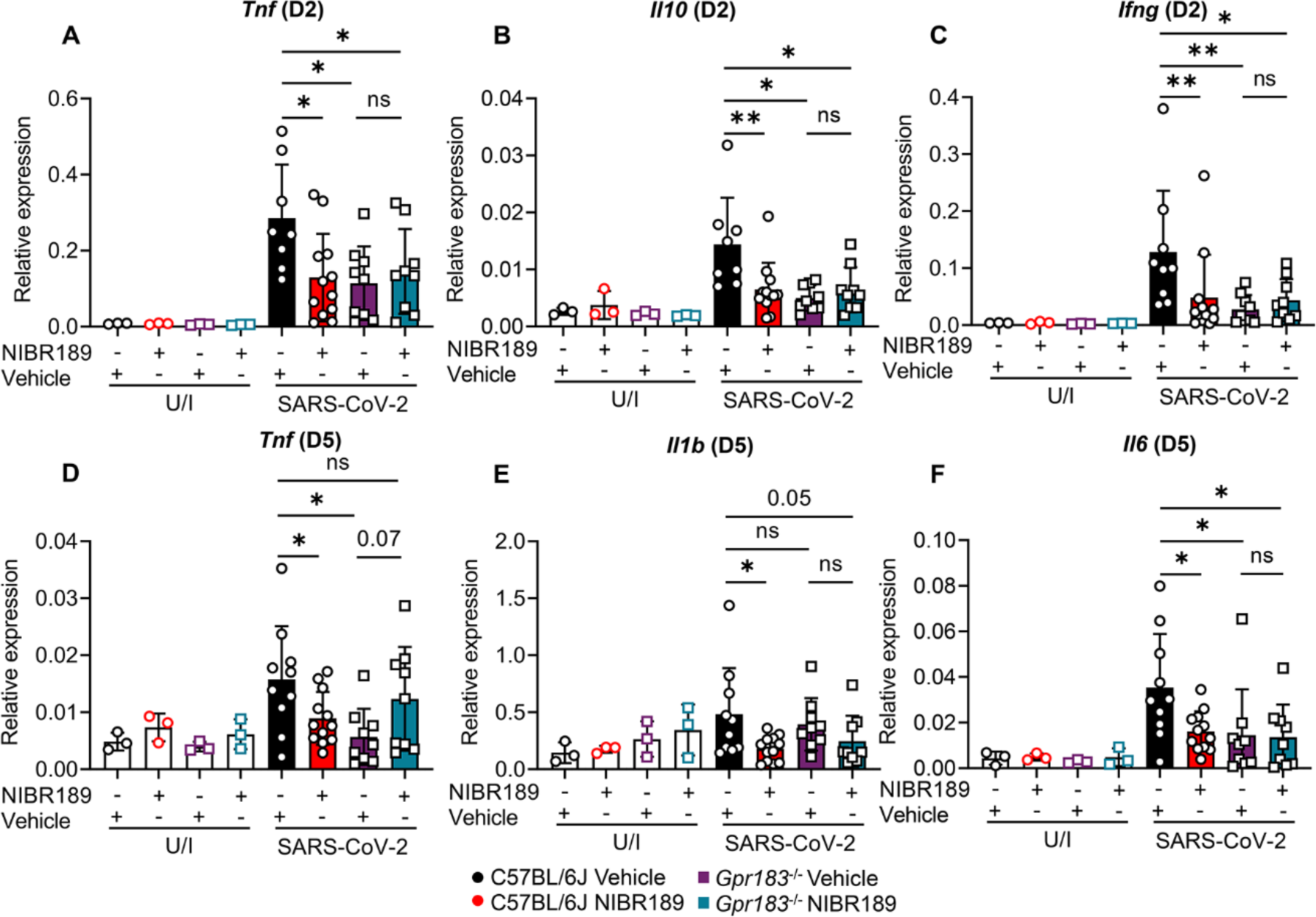
GPR183 inhibition led to reduced inflammatory cytokine profile. C57BL/6J and *Gpr183^-/-^*mice were infected intranasally with approximately 8x10^4^ PFU of mouse-adapted SARS-CoV-2. Mice were subsequently treated orally with 7.6 mg/kg NIBR189 or vehicle control twice daily from 1 dpi until the end of the experiment. Relative expression of **A**) *Tnf*, **B**) *Il10*, **C**) *Ifng* at 2 dpi and **D**) *Tnf*, **E**) *Il1b*, **F**) *Il6* at 5 dpi in the lungs measured by RT-qPCR, normalized to *Hprt*. Data are presented mean ± SD of n=3 uninfected mice and n= 9-12 infected mice per genotype and timepoint. U/I = mock infected; dpi = days post-infection; ns = not significant; *, *P* < 0.05; **, *P* < 0.01; ***, *P* < 0.001 indicate significant differences.

**Figure 8.**
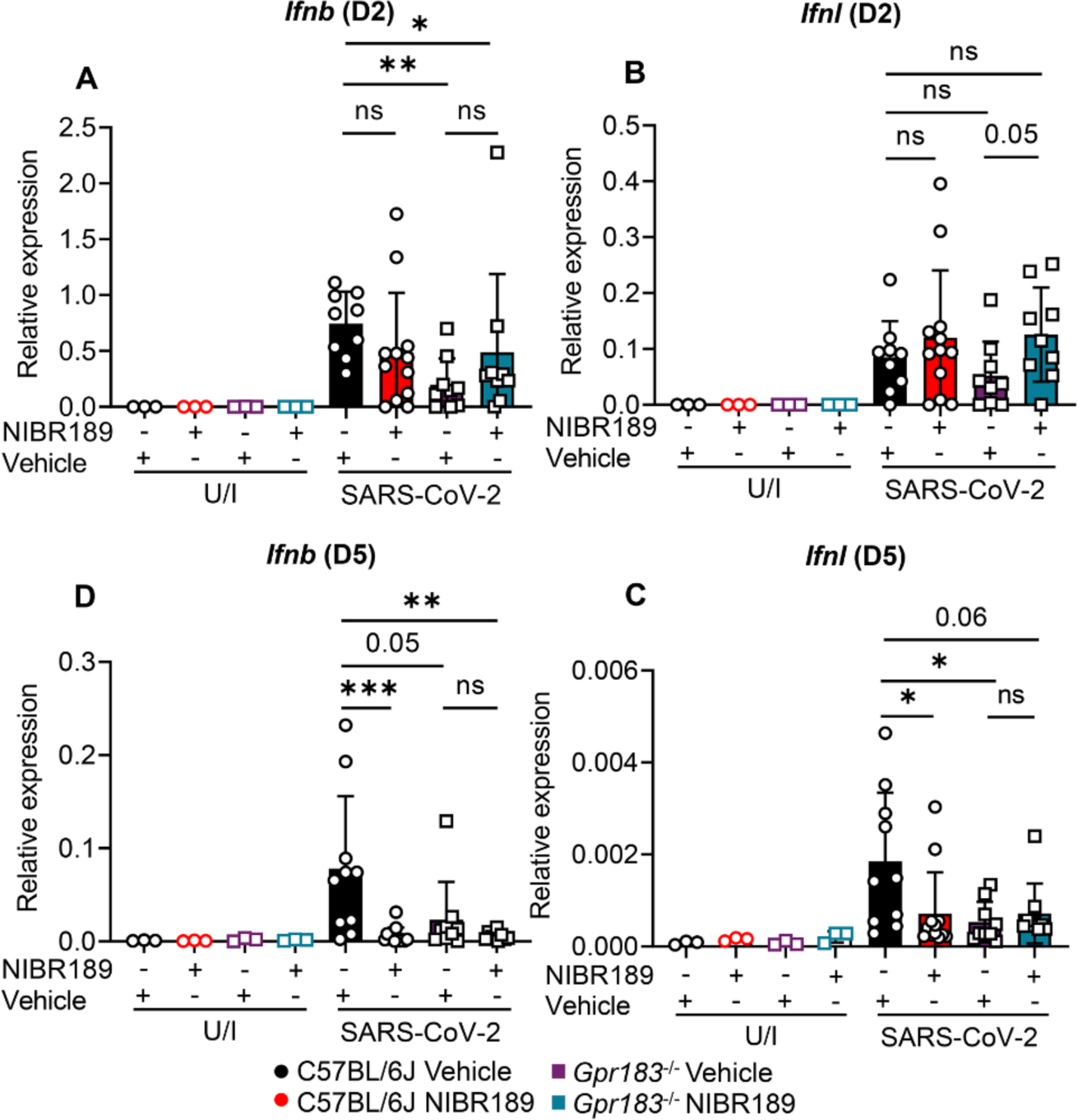
GPR183 inhibition led to reduced interferon responses at 5 dpi. C57BL/6J and *Gpr183^-/-^*mice were infected intranasally with approximately 8x10^4^ PFU of mouse-adapted SARS-CoV-2. Mice were subsequently treated orally with 7.6 mg/kg NIBR189 or vehicle control twice daily from 1 dpi until the end of the experiment. Relative expression of **A**) *Ifnb*, **B**) *Ifnl* at 2 dpi and **C**) *Ifnb*, **D**) *Ifnl* at 5 dpi in the lungs measured by RT-qPCR, normalized to *Hprt*. Data are presented mean ± SD of n=3 uninfected mice and n= 9-12 infected mice per genotype and timepoint. U/I = mock infected; dpi = days post-infection; ns = not significant; *, *P* < 0.05; **, *P* < 0.01; ***, *P* < 0.001 indicate significant differences.

### GPR183 inhibition reduces SARS-CoV-2 loads

Finally, we investigated whether the reduced macrophage infiltration and inflammatory cytokine profile in the lung of the NIBR189-treated mice is associated with altered viral loads. Viral nucleocapsid protein (Np) expression was reduced in C57BL/6J mice treated with NIBR189 compared to those administered vehicle at 2 dpi (**Figure 9A, B**). Np expression was not detected at 5 dpi, when the animals recovered from the infection. However, at the mRNA level, viral *Mpro* RNA loads in the lungs of NIBR189-treated mice were significantly lower at 5 dpi (**Figure 9C**). In summary, we demonstrate here that GPR183 inhibition reduces viral loads, macrophage infiltration and production of pro-inflammatory cytokines that are typically associated with immunopathology in the lung (**Figure 10**).

**Figure 9.**
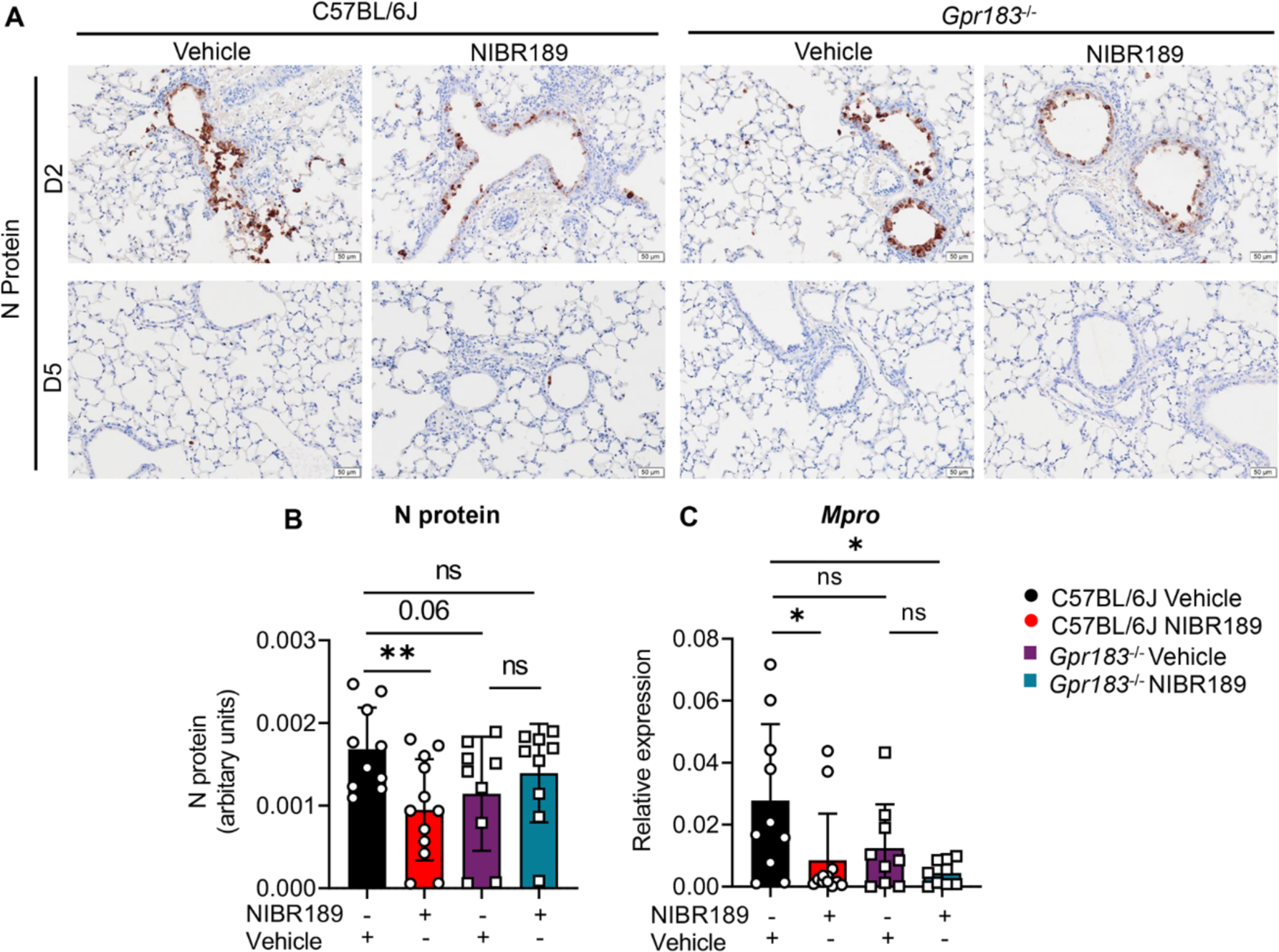
Mice treated with GPR183 antagonist had lower SARS-CoV-2 loads. C57BL/6J and *Gpr183^-/-^* mice were infected intranasally with approximately 8x10^4^ PFU of mouse-adapted SARS-CoV-2. Mice were subsequently treated orally with 7.6 mg/kg NIBR189 or vehicle control twice daily from 1 dpi until the end of the experiment. **A**) Representative IHC images of viral nucleocapsid (Np) expression at 2 dpi and 5dpi. **B**) Quantitative analysis of viral Np expression of the treatment groups at 2 dpi. **C**) Viral load was assessed in the lung through the detection of *Mpro* RNA by RT-qPCR at 5 dpi, normalized to HPRT. Data are presented mean ± SD of n=9-12 infected mice per genotype and timepoint. Scale Bar = 50μm; U/I = mock infected; dpi = days post-infection; ns = not significant; *, *P* < 0.05; **, *P* < 0.01, indicate significant differences.

**Figure 10.**
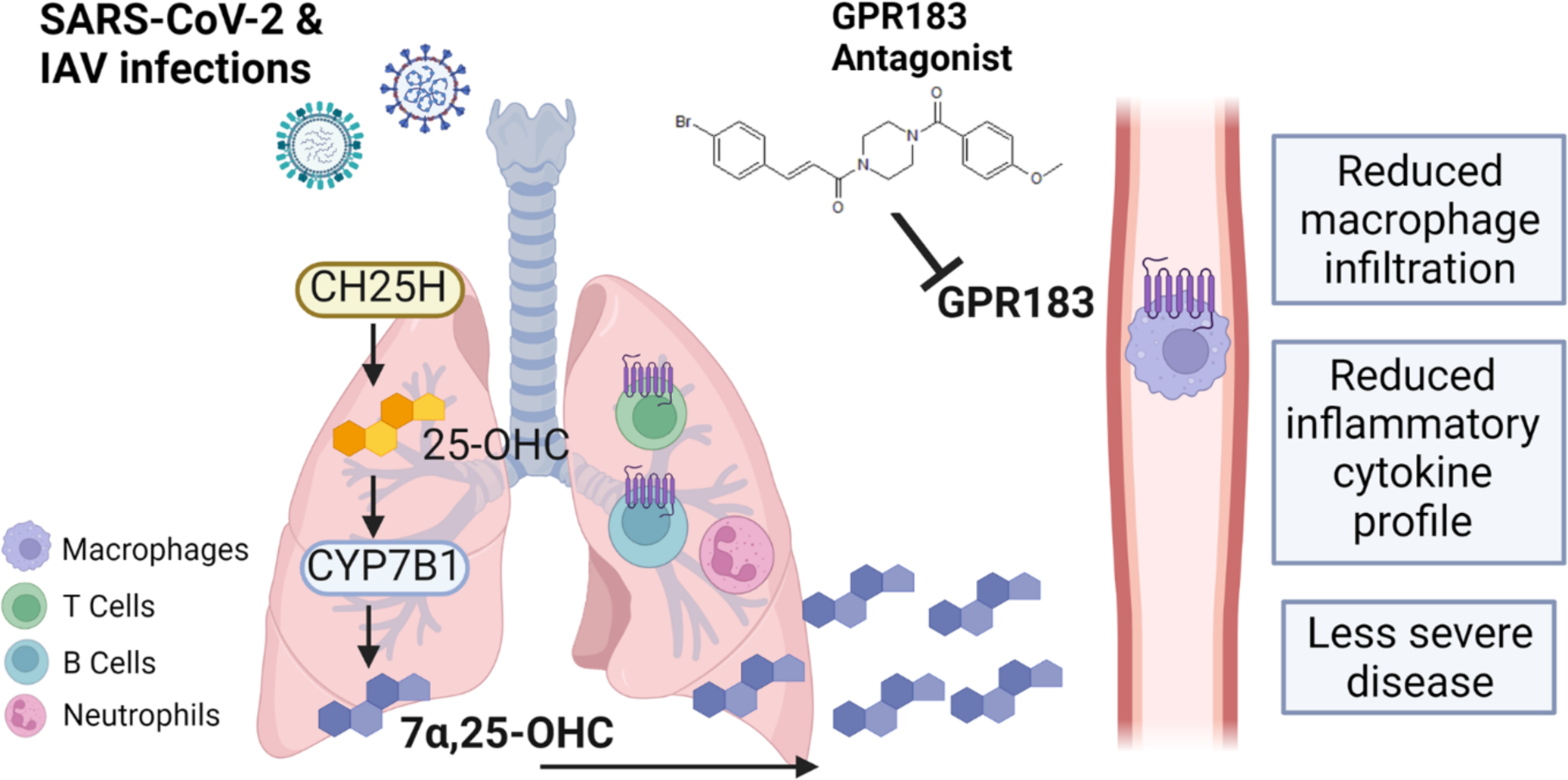
Schematic figure of the role of GPR183 in the immune response to SARS-CoV-2 and IAV infections. SARS-CoV-2 and IAV infections lead to the upregulation of CH25H and CYP7B1 which results in the production of 7α,25-OHC. This oxysterol chemotactically attracts GPR183-expressing macrophages to the lungs where they produce pro-inflammatory cytokines. Pharmacological inhibition of GPR183 attenuates the infiltration of GPR183-expressing macrophages, leading to reduced production of inflammatory cytokines without negatively affecting antiviral responses.

## Discussion

Here, we report that the oxysterols 25-OHC and 7α,25-OHC are produced in the lung upon infection with either IAV or SARS-CoV-2 and attract monocytes-macrophages in a GPR183 dependent manner to the lung. Excessive macrophage infiltration and inflammation triggers lung pathology and results in severe respiratory infection outcomes (1, 2, 29). Reduced macrophage infiltration in *Gpr183^-/-^* mice, as well as in C57BL/6J mice treated with the GPR183 antagonist NIBR189, was associated with reduced inflammatory cytokine production in the lungs of IAV and SARS-CoV-2 infected animals. Blocking GPR183 in SARS-CoV-2-infected mice significantly improved SARS-CoV-2 infection severity and attenuated viral loads. The antagonist had no impact on IAV viral loads and whether this is due to pathogen-specific effects or due to more severe disease observed by increased weight loss in the IAV model compared to the SARS-CoV-2 model, remains to be investigated. However, macrophage infiltration and cytokine production was reduced in both viral models.

In non-human primates, influenza virus infection leads to infiltration of myeloid cells into the lungs (30). Similarly, in several animal models of acute infection with SARS-CoV-2, macrophages rapidly infiltrate the lungs (4–6). Patients with severe COVID-19 infection had higher proportions of macrophages and neutrophils in BALF, with the macrophage phenotype from deceased COVID-19 patients being more activated (28). This strongly implicates macrophages as key cellular contributors to COVID-19-associated hyperinflammation. In BALF from patients with severe COVID-19, the chemokines CCL2 and CCL7 that recruit monocytes to the lung via the chemokine receptor CCR2 are also significantly enriched (31). Historically, chemokines have been considered as the main drivers of immune cell migration into the lung; however, our work here reveals that oxysterols have a non-redundant role in macrophage infiltration. Similar to our observations in *Gpr183^-/-^*mice, mice lacking the chemokine receptor CCR2 have a significant delay in macrophage infiltration into the lung (26). However, CCR2 is also required for T cell migration, therefore, animals lacking CCR2 also had delayed T cell infiltration, which correlated with significantly higher pulmonary viral titers (32). Although GPR183 is expressed on T cells it is not essential for T cell migration into the lung (33) and thus blocking GPR183 in our preclinical models did not negatively impact the T cell compartment nor other immune cell subsets.

We recently showed in a murine model of *Mycobacterium tuberculosis* (Mtb) infection that both GPR183 and CYP7B1, which produces the endogenous high affinity GPR183 agonist 7α,25-OHC, are required for rapid macrophage infiltration into the lung upon bacterial infection (24). In the Mtb model, GPR183 was also required for infiltration of eosinophils into the lung (18).

Reduced macrophage infiltration in both *Gpr183^-/-^*mice and C57BL/6J mice treated with the GPR183 antagonist NIBR189 was associated with reduced pro-inflammatory cytokine production in the lung of both IAV and SARS-CoV-2 infected animals, likely due to lower numbers of pro-inflammatory macrophages present in the tissue. However, we cannot exclude a direct effect of the GPR183 antagonist on cytokine production in macrophages and potentially other immune cell subsets like T cells. We previously showed that GPR183 is a constitutively negative regulator of type I IFNs in primary human monocytes infected with Mtb (34). *In vitro* activation of GPR183 with the agonist 7α,25-OHC reduced Mtb-induced *Ifnb* mRNA levels, while the GPR183 antagonist GSK682753 significantly increased *Ifnb* mRNA expression elicited by Mtb (34). This antagonist did not affect *Tnf* transcription in these *in vitro* assays; however, it cannot be excluded that NIBR189 used in the experiments presented here directly affects cytokine expression in macrophages or other immune cell subsets.

Irrespective of the exact mechanism, reduced pro-inflammatory cytokine production was associated with reduced SARS-CoV-2 infection severity. Excessive production of proinflammatory cytokines contributes to the immunopathology in COVID-19 patients with severe disease (35). Therefore, lower pro-inflammatory cytokine production in animals treated with NIBR189 can explain, at least in part, the better disease outcomes compared to vehicle-treated animals. While cytokines can be detrimental to the host and contribute to the development of cytokine storms (36), early type I and III IFNs are crucial in controlling viral replication during IAV (37, 38) and SARS-CoV-2 infections (39, 40), whereas prolonged type I IFN responses can be detrimental to the host (41). The GPR183 antagonist did not alter early type I or III IFN responses in SARS-CoV-2-infected animals, suggesting that the anti-viral response was not impaired by the treatment. However, antagonising GPR183 prevented a prolonged IFN response, which was associated with more effective viral clearance observed in NIBR189-treated animals.

While several oxysterols can have a direct anti-viral effect (12), it is not known whether NIBR189 directly affects viral entry or replication. CH25H/25-OHC have been shown to inhibit SARS-CoV-2 infection *in vitro* by blocking the virus-host cell membrane fusion (42, 43). It is unlikely that NIBR189 directly affects viral entry and/or replication, given that it is structurally very different from cholesterols and probably not able to disrupt the host cell membrane composition typical for other anti-viral oxysterols.

We propose that GPR183, which belongs to the GPCR family, is a novel drug target for severe COVID-19. GPCRs are popular targets because of their pharmacological tractability. Indeed, 34% of all FDA approved drugs are directed against members of this receptor family, with this accounting for global sales volumes of over 180 billion US dollars (44). In our SARS-CoV-2 model the GPR183 antagonist demonstrated a dual benefit by not only reducing pro- inflammatory cytokines without compromising early type I and type III IFN responses, but also by reducing viral loads. Other immunosuppressive therapies used in severe COVID-19 like glucocorticoids can increase ACE2 expression which promotes viral entry and replication (45, 46). Consistent with this, glucocorticoid use delays SARS-CoV-2 clearance (47). Glucocorticoids can also affect antibody production. While it remains to be established whether NIBR189 has a similar effect, short term use of a GPR183 antagonist during the acute viral infection is unlikely to negatively impact antibody responses. Currently available antiviral treatments are effective, but mutations in SARS-CoV-2 conferring resistance to new antivirals are already emerging (48). Therefore, adjunct host-directed therapy with a GPR183 antagonist together with conventional antivirals may increase treatment efficacy. Since a GPR183 antagonist targets the host and not the virus it is not anticipated that viruses will develop resistance against host directed therapy (49). Further, a GPR183 antagonist-based therapy can also be immediately effective against newly emerging SARS-CoV-2 variants without further adaption.

In summary, we provide the first preclinical evidence of GPR183 as a novel host target for therapeutic intervention to reduce macrophage-mediated hyperinflammation, SARS-CoV-2 loads and disease severity in COVID-19.

## Acknowledgements

This study was supported by grants to KR from the Mater Foundation, the Australian Respiratory Council, Diabetes Australia, and the Australian Infectious Diseases Research Centre. SB was supported by an early career seed grant from the Mater Foundation. The Translational Research Institute is supported by a grant from the Australian Government. We thank A/Prof Sumaira Hasnain for sharing antibodies used in this study. We thank the Queensland Health Forensic and Scientific Services, Queensland Department of Health, for providing SARS-CoV-2 isolate. We acknowledge the technical assistance of the team that operates and maintains the Australian Galaxy service (https://usegalaxy.org.au/). The Danish Council for Independent Research I Medical Sciences supported MMR. MJS is supported by a National Health and Medical Research Council of Australia Investigator grant (APP1194406). KRS is funded by the NHMRC Investigator Grant (2007919) and is consultant for Sanofi, Roche and NovoNordisk. MMR is co-founder of Antag Therapeutics and of Synklino. The opinions and data presented in this manuscript are of the authors and are independent of these relationships. Other authors declare no competing interests. We thank Profs David Hume, Jean-Pierre Levesque and Maher Gandhi for critical review of the manuscript.

## Author contributions

Conceptualization: CXF, SB, MJS, KRS, MMR, KR Methodology: KYC, HBO, BJA, BM,SR Investigation: CXF, SB, KYC, MDN, HBO, BJA, BM, SR, RW, LB, JES, RP, AK Writing-original draft: CXF, SB, KR Writing-review and editing: all authors. Funding acquisition: SB, KRS, MMR, KR.

---

**Figure S1.**
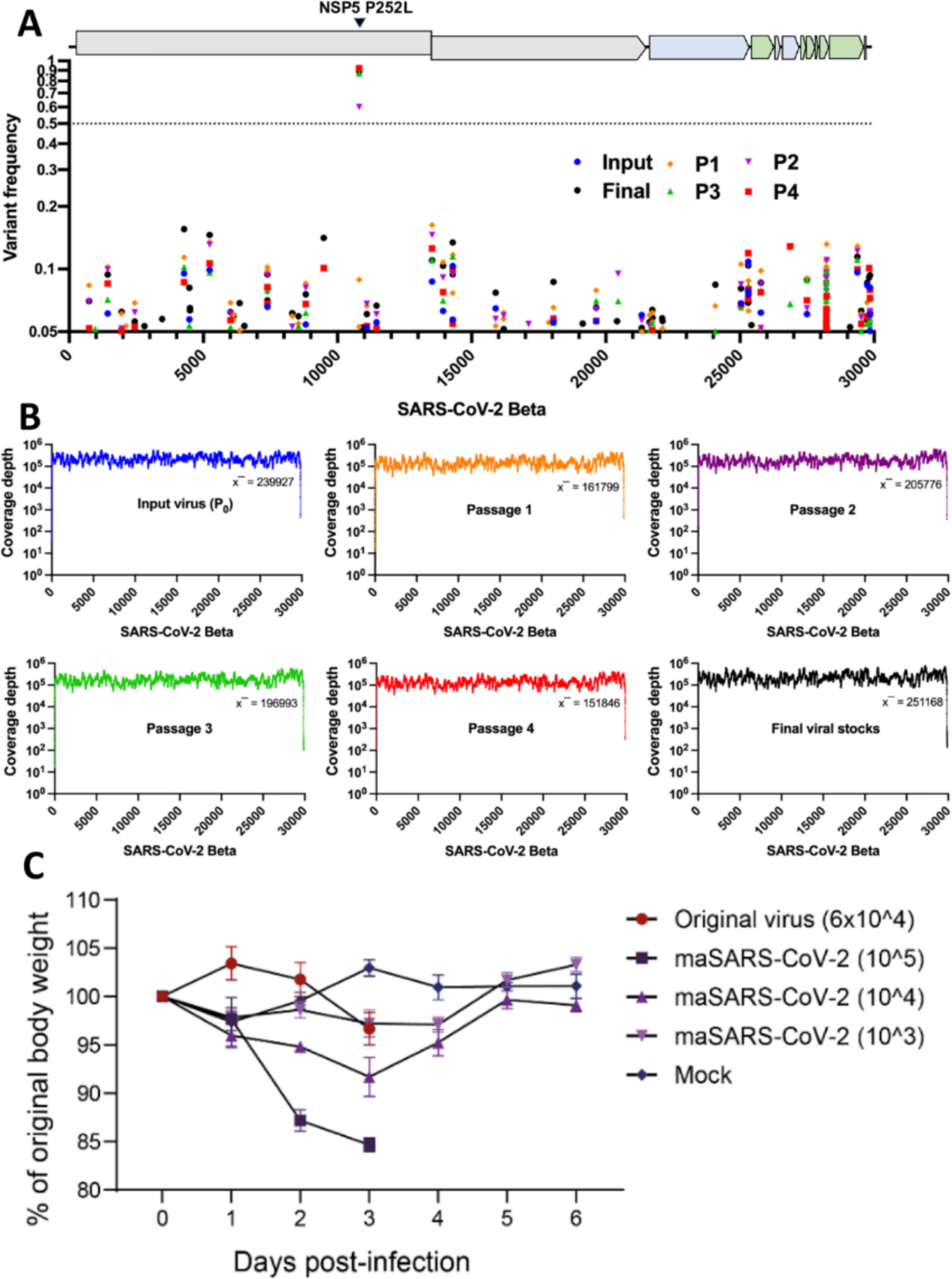
Evolution and coverage of mouse-adapted SARS-CoV-2 virus. (**A**) Mutation frequency of input SARS-CoV-2 Beta virus (blue circle) and passage one (orange diamond), passage two (purple nabla), passage three (green triangle) and passage four (red square) mouse-adapted viruses over the reference genome sequence as well as the final virus stocks (black circle) amplified in VeroE6-hTMPRSS2 cells. The dotted line indicates the consensus frequency of 0.5 **B**) Summary plots of read coverage of passaged SARS-CoV-2 viruses from A) mapping to SARS-CoV-2 Beta strain. Depth of coverage of binary alignment files was determined using samtools depth. **C**) Weight loss over time following infection with the Beta variant of SARS-CoV-2 (original virus) or various doses of maSARS-CoV-2 (after four passages in mice). Plaque forming units are indicated in brackets. Data indicates mean ± SEM.

**Figure S2.**
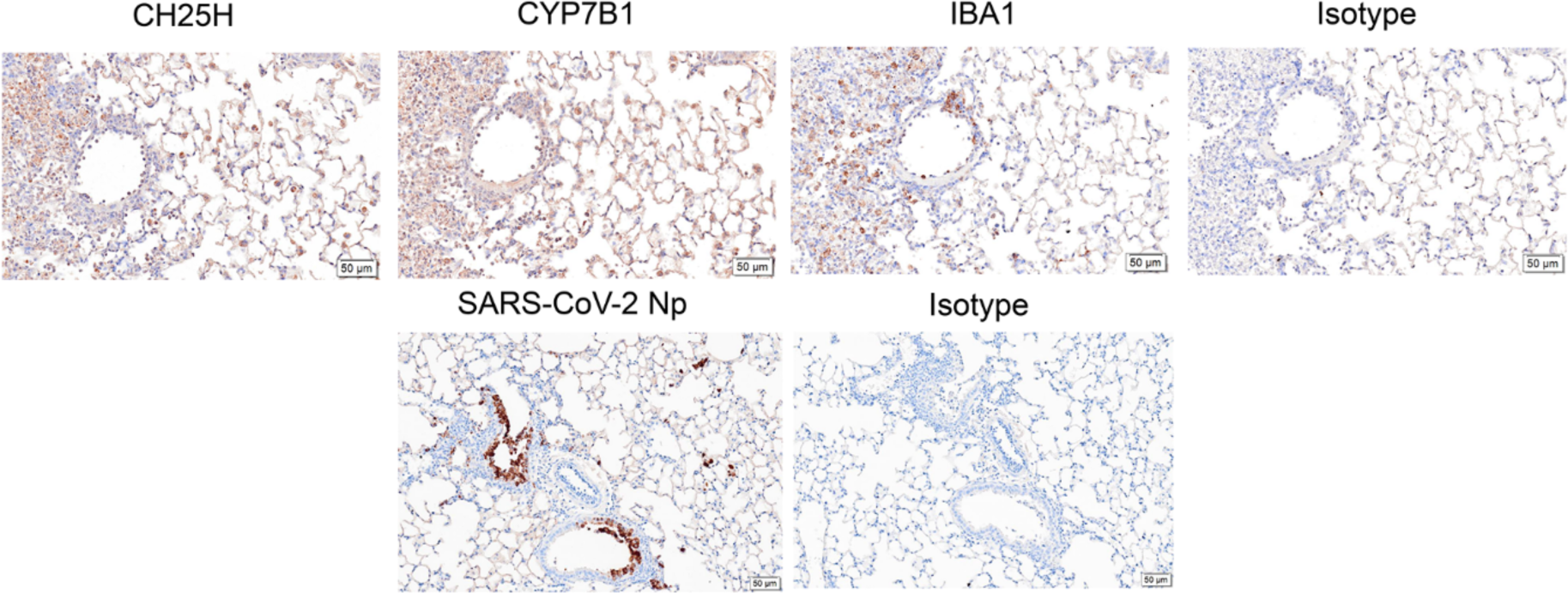
Isotype staining controls for CYP7B1, CH25H, IBA1 and viral Np. IHC of IAV-infected lung sections incubated with rabbit anti-CH25H, rabbit anti-CYP7B, rabbit anti-IBA1 and an isotype-matched control (Rabbit IgG; negative control). IHC of SARS-CoV-2-infected lung sections incubated with rabbit anti-SARS-CoV-2 nucleocapsid protein (Np) and an isotype-matched control (Rabbit IgG; negative control). Scale bar = 50μm

**Figure S3.**
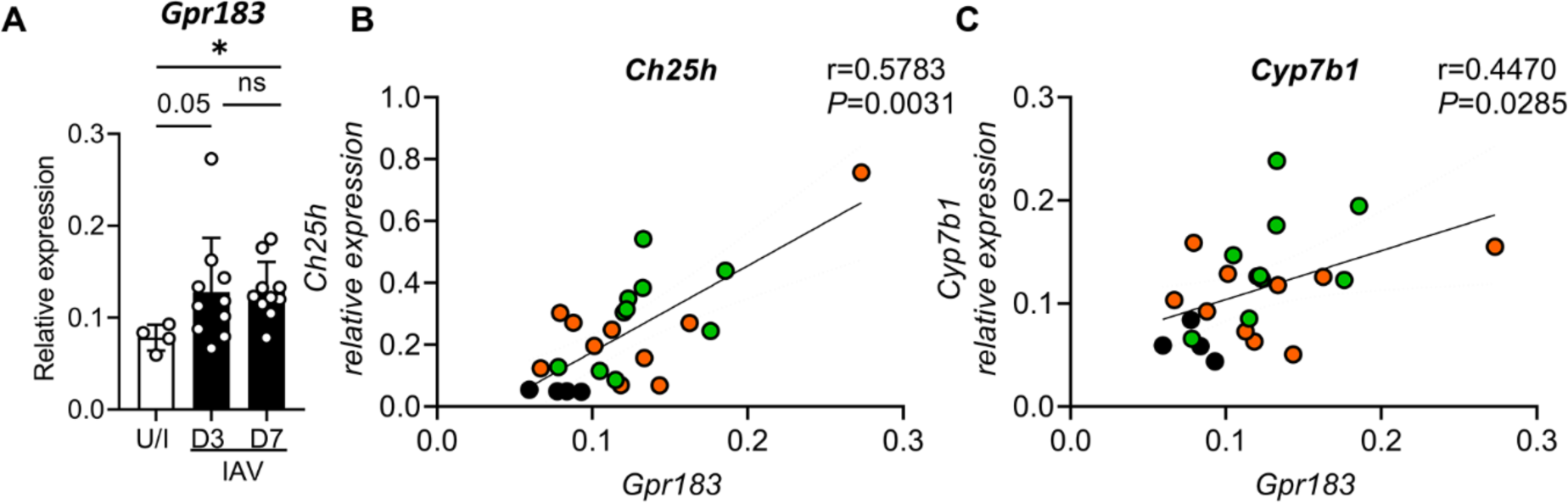
*Gpr183* mRNA expression is upregulated in the lung during IAV infection and correlates with expression of the oxysterol synthesising enzymes CH25H and CYP7B1. C57BL/6J mice were infected intranasally with 5,500 PFU of A/Auckland/01/09. **A**) Relative expression of *Gpr183* mRNA measured by RT-qPCR, normalized to *Hprt*. Correlation analyses were performed with mRNA expression levels of *Gpr183* and oxysterol synthesizing enzymes. Individual scatter plots showing correlations between *Gpr183* and **B**) *Ch25h* and **C**) *Cyp7b1*. Black dots represent uninfected samples while coloured dots represent IAV-infected samples (Orange dots, 3 dpi; green dots, 7 dpi). Data are presented as mean ± SD of n=4 uninfected and n=8-10 infected mice per timepoint. ns = not significant; *, *P* < 0.05 indicate significant differences. Spearman rank correlation test were used to calculate correlation coefficient and to determine significant correlations with values displayed on each scatter plot.

**Figure S4.**
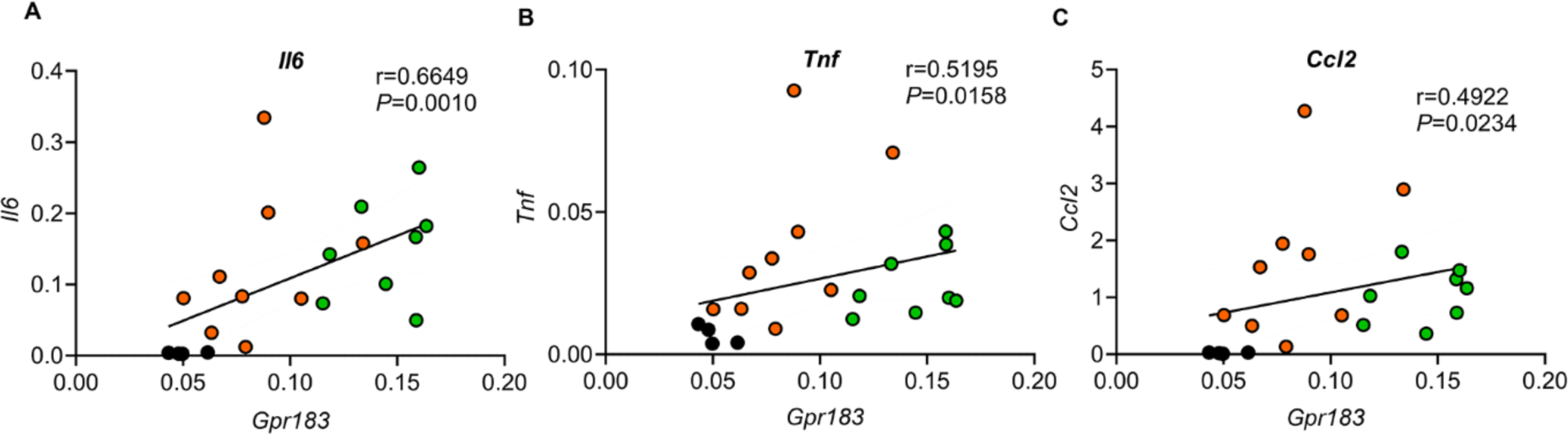
Correlations between lung mRNA expression of *Gpr183* and inflammatory markers in IAV-infected mice. Correlation analyses of *Gpr183* mRNA expression with mRNA expression of inflammatory cytokines in lung tissue from IAV-infected C57BL/6J mice (n=21 pairs). Relative gene expression was determined by RT-qPCR, normalized to *Hprt*. Individual scatter plots showing correlations between *Gpr183* and **A**) *Il6*, **B**) *Tnf* and **C**) *Ccl2*. Black dots represent uninfected samples while coloured dots represent IAV-infected samples (Orange dots, 3dpi; green dots, 7dpi). Spearman rank correlation test were used to calculate correlation coefficient and to determine significant correlations with values displayed on each scatter plot.

**Figure S5.**
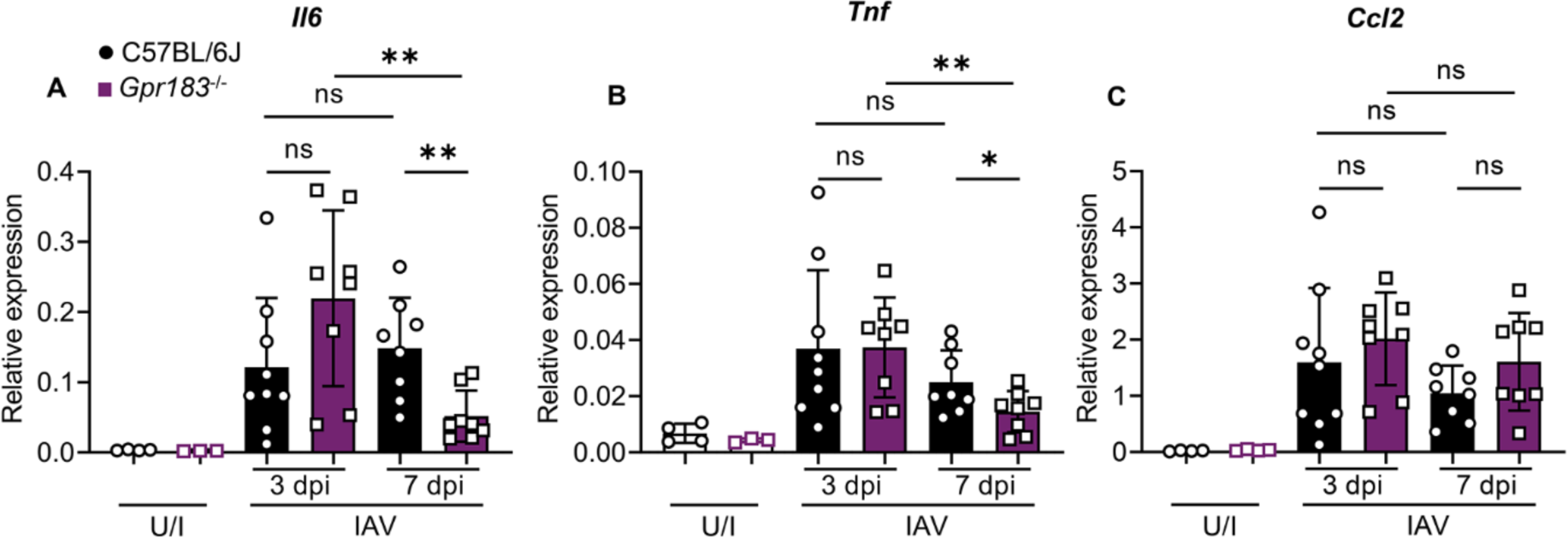
Cytokine expression at mRNA and protein level in IAV-infected C57BL/6J and *Gpr183^-/-^* mice. C57BL/6J and *Gpr183^-/-^* mice were infected intranasally with 5,500 PFU of A/Auckland/01/09. Cytokine measurements of **A**) *Il6,* **B**) *Tnf* and **C**) *Ccl2* at 3 dpi and 7 dpi measured by RT-qPCR, normalized to *Hprt.* Data are presented as mean ± SD of n=4 uninfected per genotype and n=8-10 infected mice per genotype and timepoint. U/I = uninfected; dpi = days post-infection; ns = not significant; *, *P* < 0.05; **, *P* < 0.01 indicate significant differences.

**Figure S6.**
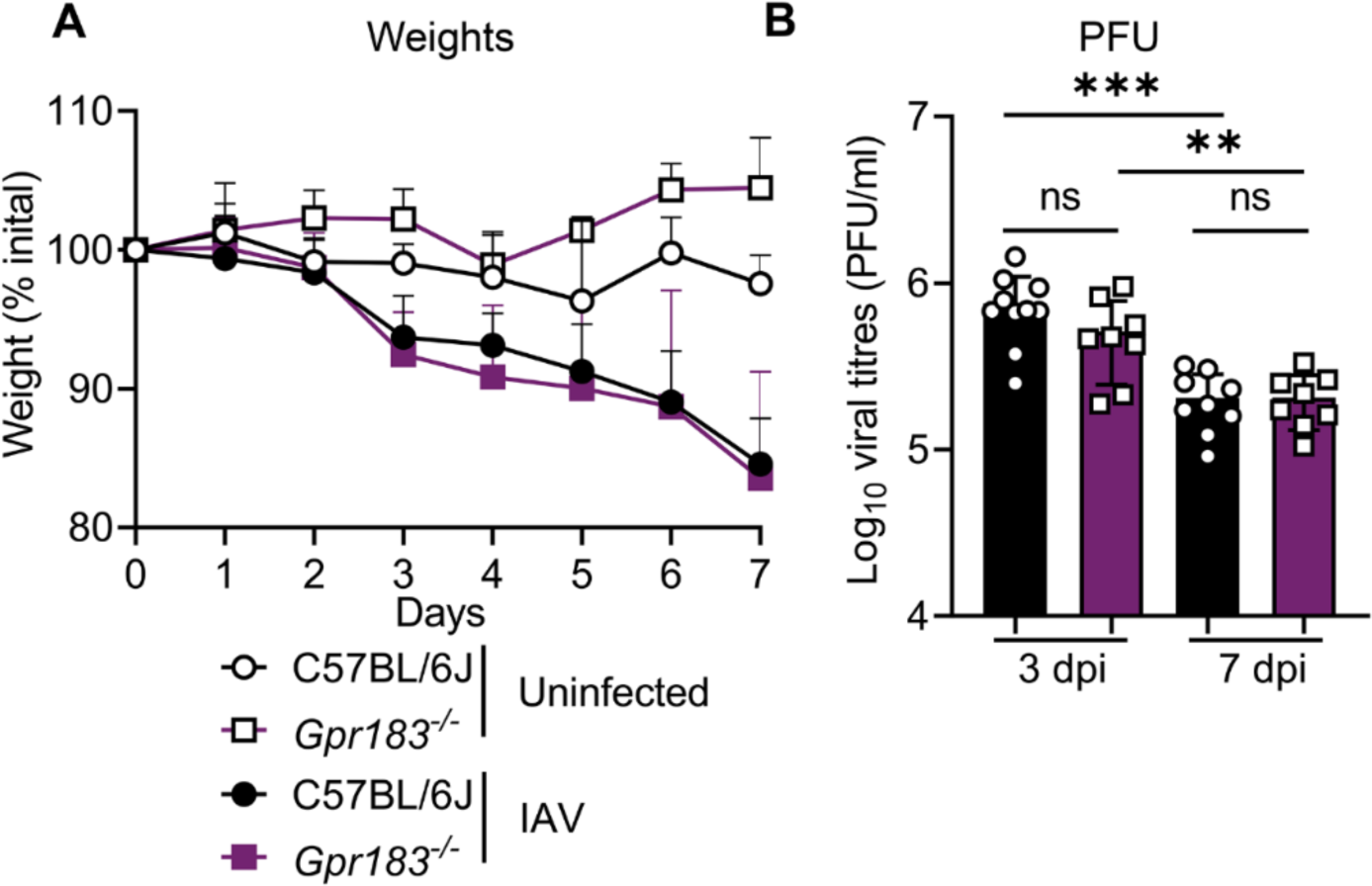
Weights of IAV and mock infected C57BL/6J and *Gpr183^-/-^*mice and viral loads. C57BL/6J and *Gpr183^-/-^* mice were infected intranasally with approximately 5,500 PFU of A/Auckland/01/09. **A**) Weights of IAV- or mock-inoculated mice are displayed as percentage of the weight at time of inoculation. **B**) Viral load was assessed by measuring the PFU by plaque assays. Data are presented as mean ± SD for n=8-10 infected mice per genotype and timepoint. dpi = days post-infection; ns = not significant; **, *P* < 0.01; ***, *P* < 0.001 indicate significant differences.

**Figure S7.**
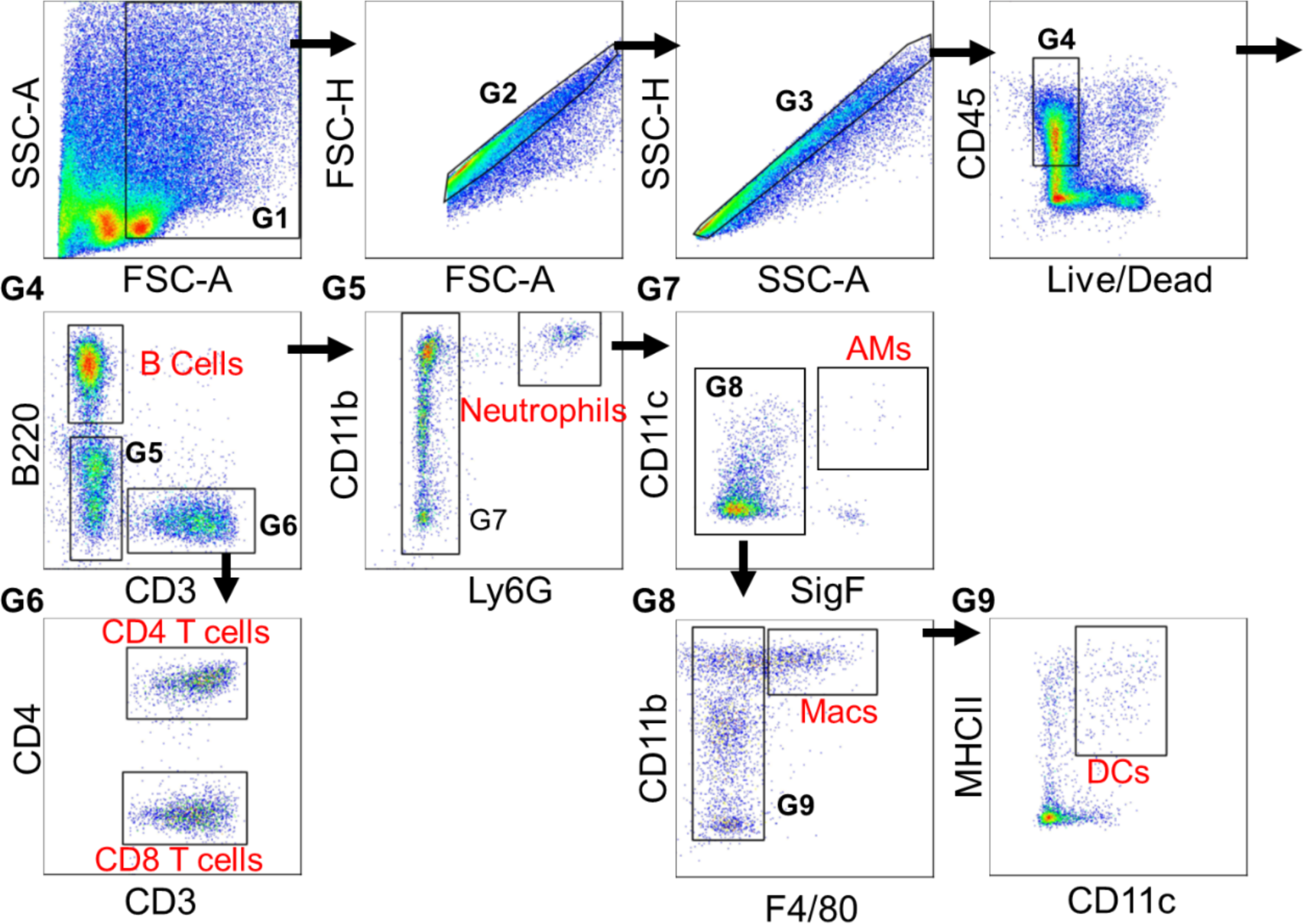
Representative flow cytometry plots illustrating the gating strategy of immune cells. C57BL/6J or *Gpr183*^-/-^ mice were infected intranasally with 5,500 PFU of A/Auckland/01/09. Mice were subsequently treated orally with 7.6 mg/kg NIBR189 or vehicle control twice daily from 1 dpi until the end of the experiment. Gates containing multiple cell populations are numbered (G1-G9). Gates that contained a single cell population are labeled with its respective cell type. These includes: B cells (B220^+^; G5), CD4^+^ T cells (CD3^+^,CD4^+^; G6), CD8^+^ T Cells (CD3^+^,CD4^-^; G6), Neutrophils (B220^-^,CD3^-^,Ly6G^+^; G5), Alveolar macrophages (B220^-^,CD3^-^,Ly6G^-^,CD11c^+^,SigF^+^; G7), Macrophages (B220^-^,CD3^-^,Ly6G^-^,SigF^-^,CD11b^+^,F4/80^high^; G8) and Dendritic cells (DCs; B220^-^,CD3^-^,Ly6G^-^,SigF^-^, F4/80^low^,CD11c^+^, MHCII^+^; G9).

**Figure S8.**
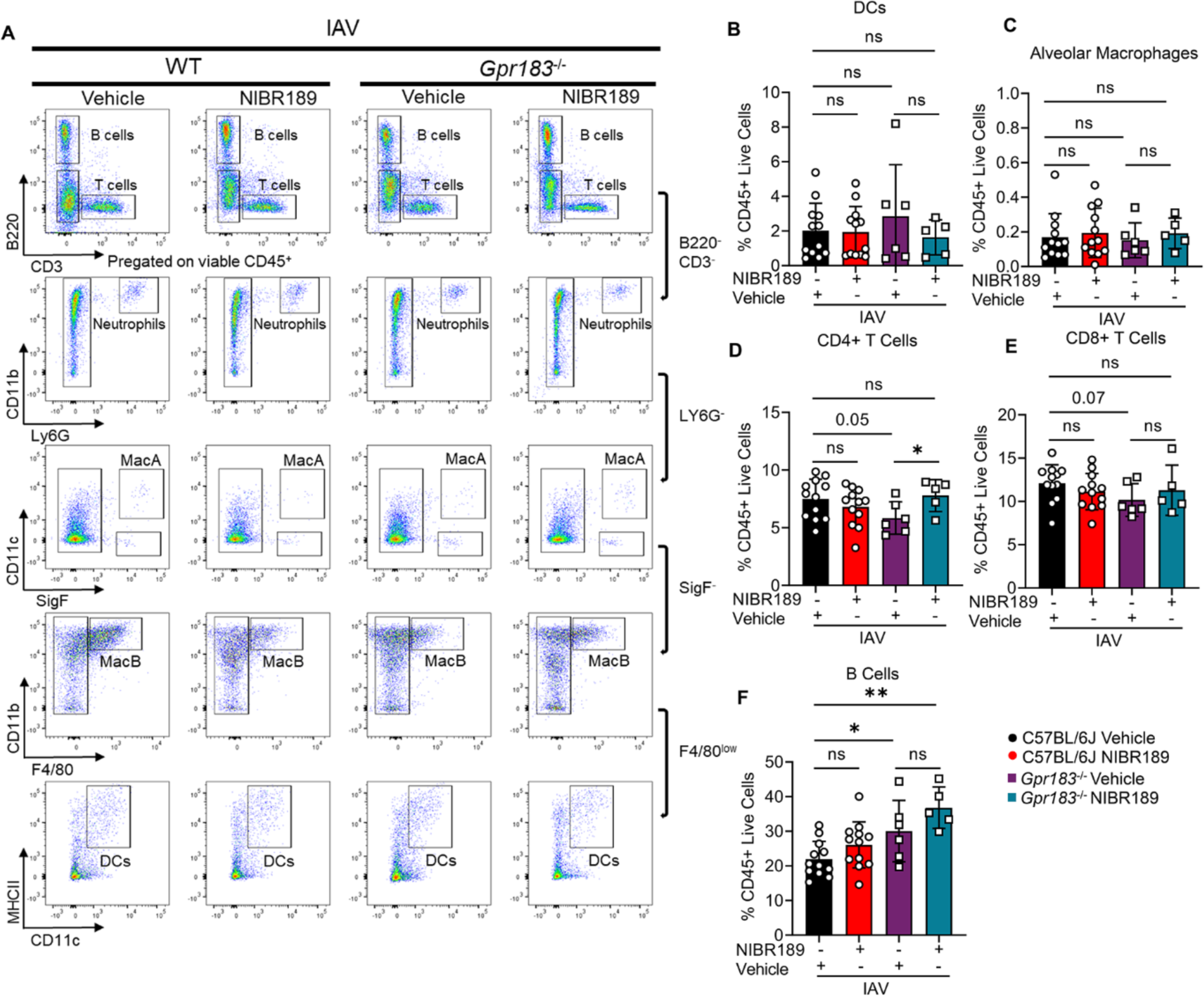
Immune cell populations in the lungs of IAV-infected mice treated with the GPR183 antagonist NIBR189. C57BL/6J or *Gpr183^-/-^*mice were infected intranasally with 5,500 PFU of A/Auckland/01/09. Mice were subsequently treated orally with 7.6 mg/kg NIBR189 or vehicle control twice daily from 1 dpi until the end of the experiment. **A**) Frequency of B cells (B220^+^), T cells (CD3^+^ CD8^+^ or CD4^+^), neutrophils (B220^-^CD3^-^Ly6G^+^) was determined by flow cytometry against total viable CD45^+^ immune cells at 3 dpi. Alveolar macrophages (CD11c^+^SigF^+^), infiltrating macrophages (F480^high^/CD11b^+^/Ly6G^-^/SigF^-^) and dendritic cells (SigF^-^F4/80^-^MHCII^+^CD11c^+^) were further identified from the B220^-^CD3^-^Ly6G^-^ cell population. (**B-G**) Graphs depicting the frequency of **B**) Dendritic cells, **C**) alveolar macrophages, **D**) CD4^+^ T cells, **E**) CD8^+^ T cells and **F**) B cells against total viable CD45^+^ immune cells. Data are presented mean ± SD of n=5-12 infected mice per genotype and timepoint. UI = uninfected; dpi = days post-infection; ns = not significant.

**Figure S9.**
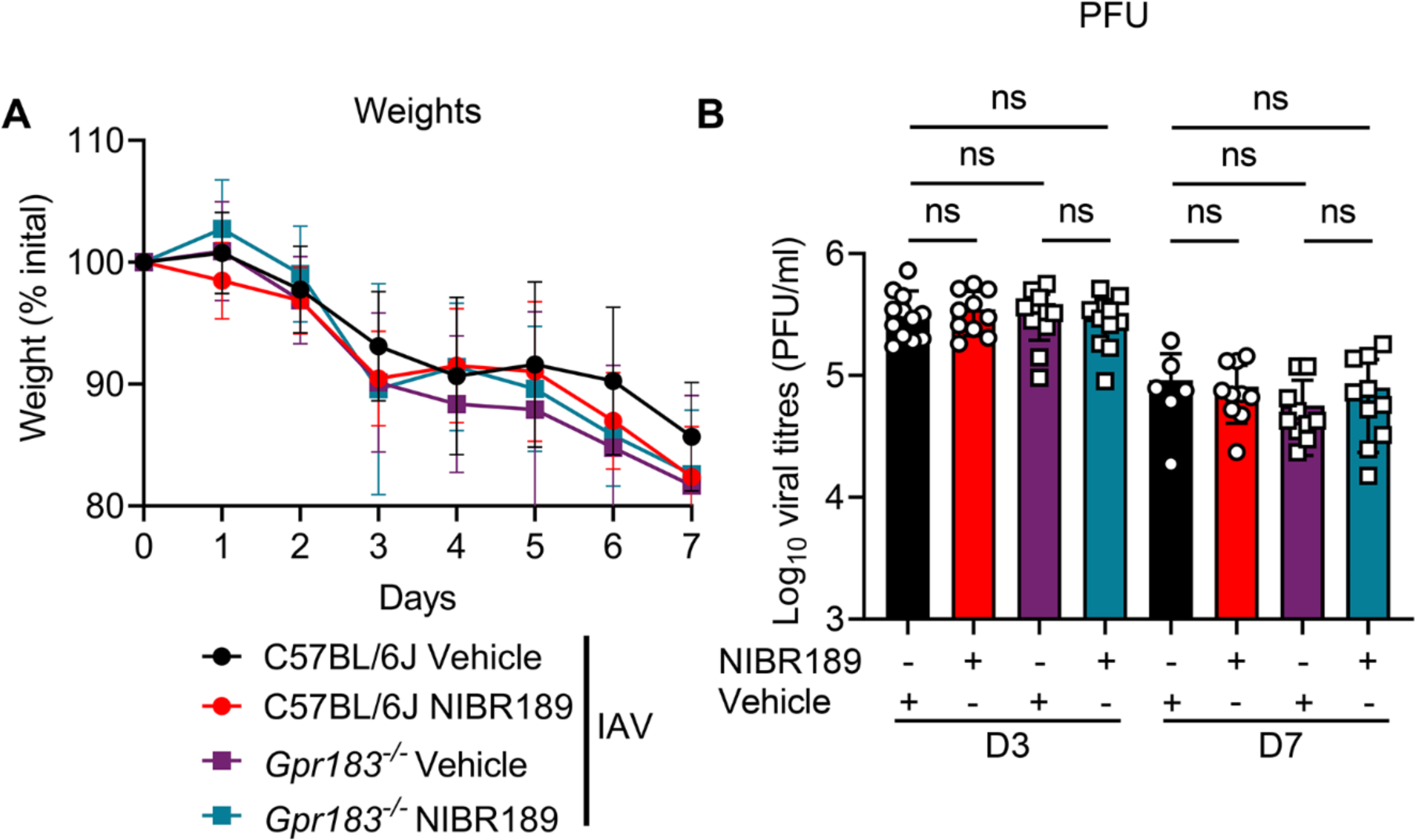
Body weights and viral loads of IAV-infected C57BL/6J and *Gpr183^-/-^*mice treated with NIBR189 or vehicle. C57BL/6J mice and *Gpr183^-/-^* mice were infected intranasally with 5,500 PFU of A/Auckland/01/09. Mice were subsequently treated orally with 7.6 mg/kg NIBR189 or vehicle control twice daily from 1 dpi until the end of the experiment. **A**) Weights of IAV- or mock-inoculated mice with or without treatment are displayed as percentage of the weight at time of inoculation. **B**) Viral load was assessed by measuring the PFU through plaque assay. Data are presented mean ± SD of n=6-12 infected mice per genotype and timepoint. UI = uninfected; dpi = days post-infection; ns = not significant.

**Figure S10.**
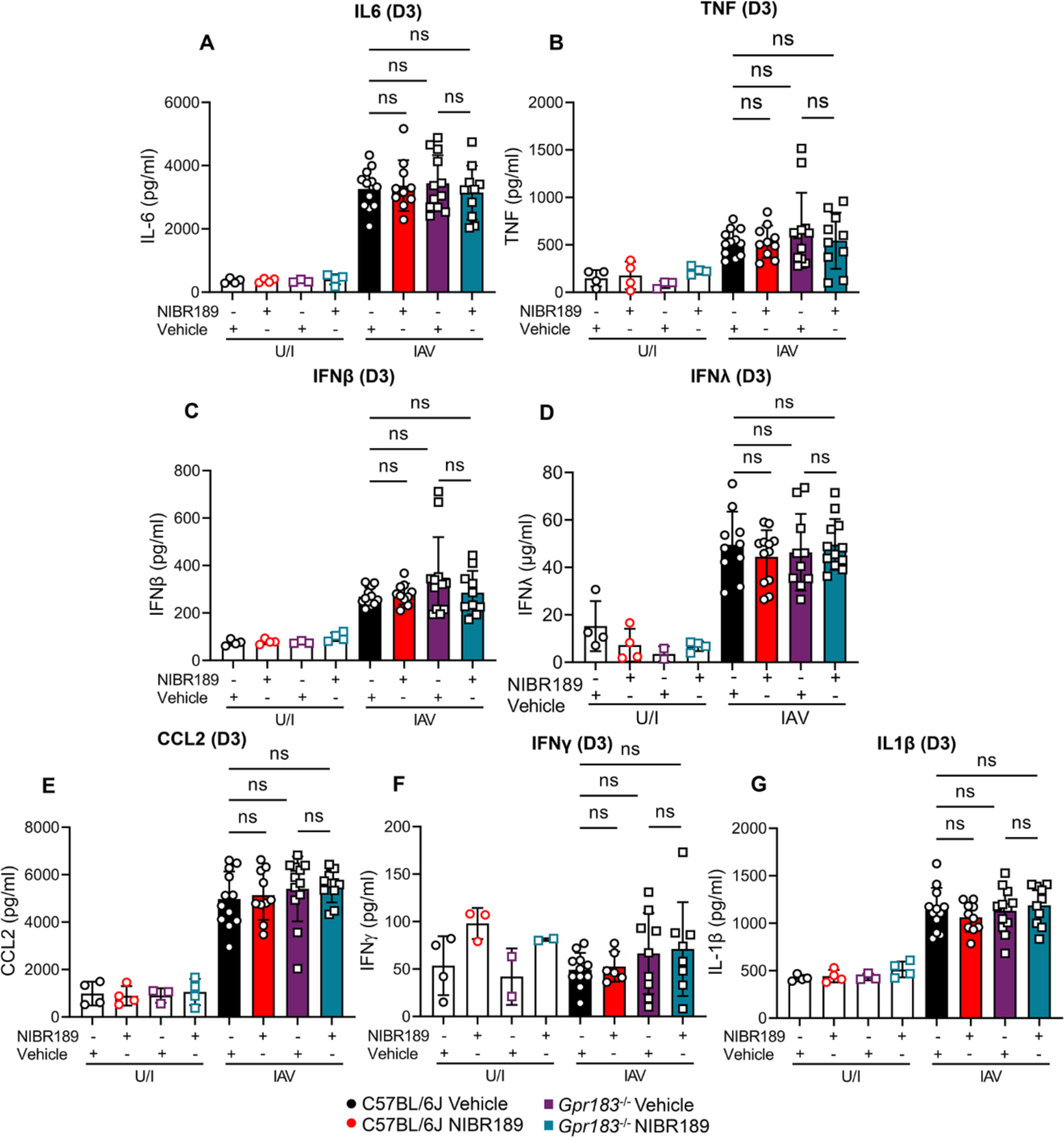
Cytokine expression at protein level in IAV-infected C57BL/6J and *Gpr183^-^* *^/-^* mice treated with NIBR189 and/or vehicle. C57BL/6J and *Gpr183^-/-^*mice were infected intranasally with 5,500 PFU of A/Auckland/01/09. Mice were subsequently treated orally with 7.6 mg/kg NIBR189 or vehicle control twice daily from 1 dpi until the end of the experiment. Cytokine measurements of **A**) IL-6 **B**) TNF, **C**) IFNβ, **D**) IFN*λ*, **E**) CCL2, **F**) IFNγ and **G**) IL-1β, at 3 dpi measured by ELISA. Data are presented mean ± SD of n=4 uninfected mice per genotype and n=6-12 infected mice per genotype. U/I = uninfected; dpi = days post-infection; ns = not significant. *, *P* < 0.05 indicate significant differences.

**Figure S11.**
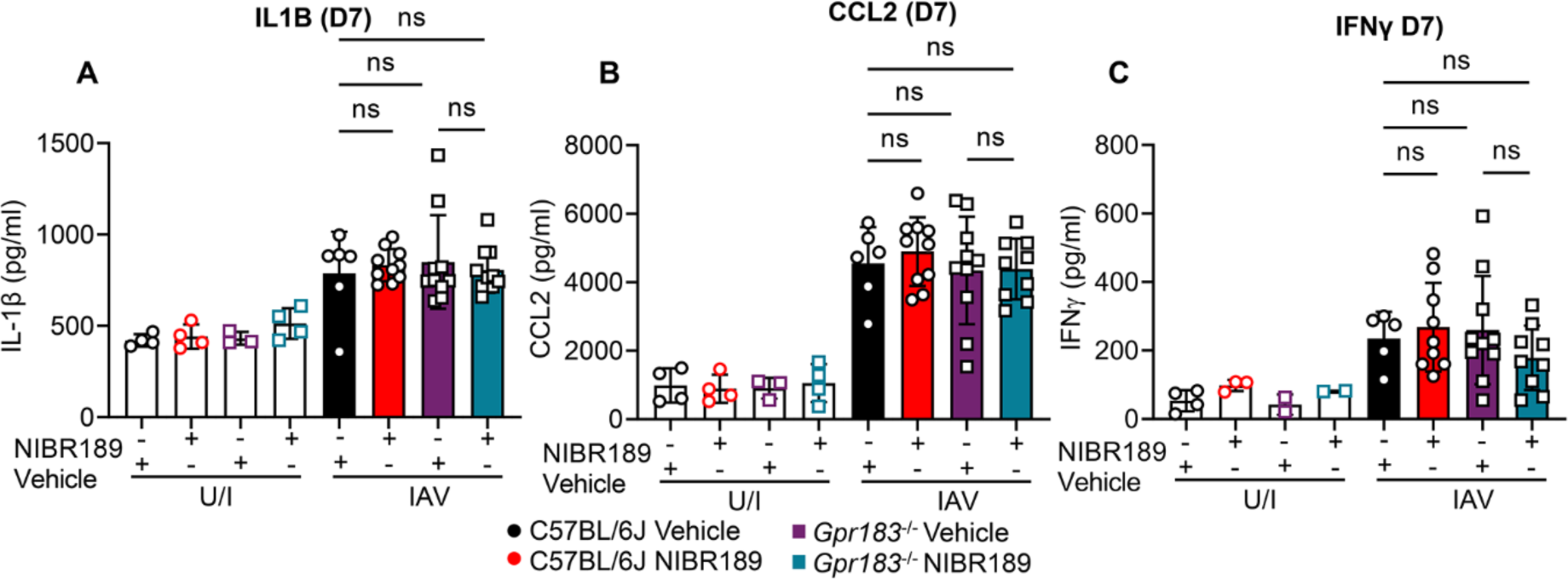
Cytokine expression at protein level in IAV-infected C57BL/6J and *Gpr183^-^* *^/-^* mice treated with NIBR189 and/or vehicle. C57BL/6J and *Gpr183^-/-^* mice were infected intranasally with 5,500 PFU of A/Auckland/01/09. Mice were subsequently treated orally with 7.6 mg/kg NIBR189 or vehicle control twice daily from 1 dpi until the end of the experiment. Cytokine measurements of **A**) IL-1β, **B**) CCL2, and **C**) IFNγ at 7 dpi measured by ELISA. Data are presented mean ± SD of n=4 uninfected mice per genotype and n=6-12 infected mice per genotype. U/I = uninfected; dpi = days post-infection; ns = not significant.

**Figure S12.**
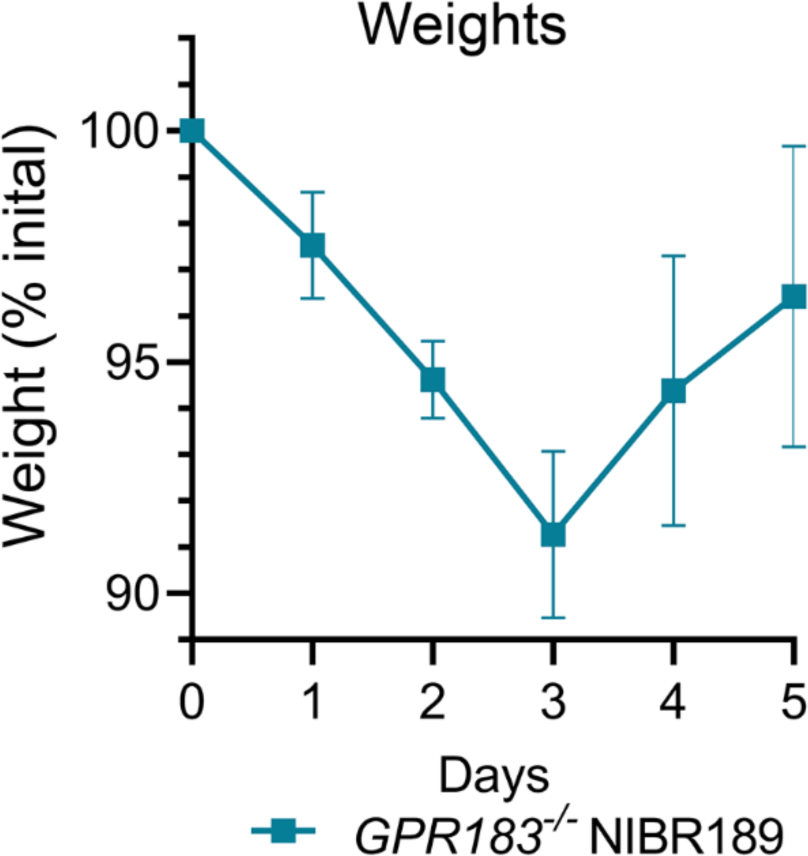
GPR183 inhibition weight loss upon SARS-CoV-2 infection. C57BL/6J and *Gpr183^-/-^* mice were infected intranasally with approximately 8x10^4^ PFU of mouse-adapted SARS-CoV-2. Mice were subsequently treated orally with 7.6 mg/kg NIBR189 or vehicle control twice daily from 1 dpi until the end of the experiment. Weights of mice displayed as percentage of the weight at time of inoculation.

**Figure S13.**
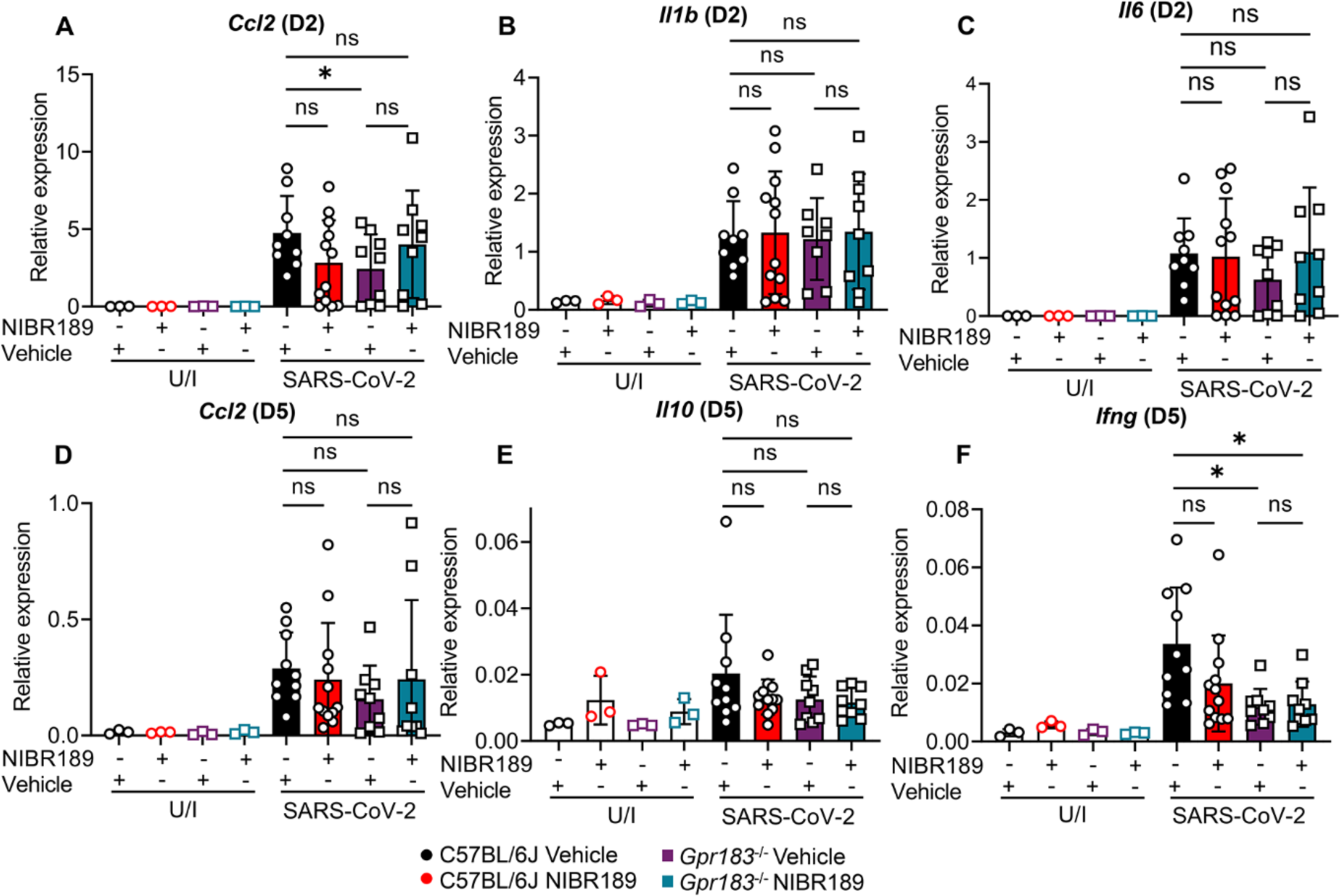
Cytokine expression at mRNA in SARS-CoV-2-infected C57BL/6J and *Gpr183^-/-^* mice treated with GPR183 antagonist at 2 dpi and 5 dpi. C57BL/6J and *Gpr183^- /-^* mice were infected intranasally with approximately 8x10^4^ PFU of mouse-adapted SARS-CoV-2. Mice were subsequently treated orally with 7.6 mg/kg NIBR189 or vehicle control twice daily from 1 dpi until the end of the experiment. Expression of **A**) *Ccl2*, **B**) *Il1b* and **C**) *Il6* at 2 dpi and **D**) *Ccl2*, **E**) *Il10* and **F**) *Ifng* 5 dpi was measured by RT-qPCR normalized to HPRT. Data are presented mean ± SD of n=3 uninfected mice and n= 9-12 infected; mice per genotype and timepoint. U/I = uninfected dpi = days post-infection; ns = not significant. *, *P* < 0.05; **, *P* < 0.01; ***, *P* < 0.001 indicate significant differences.

**Table S1:**
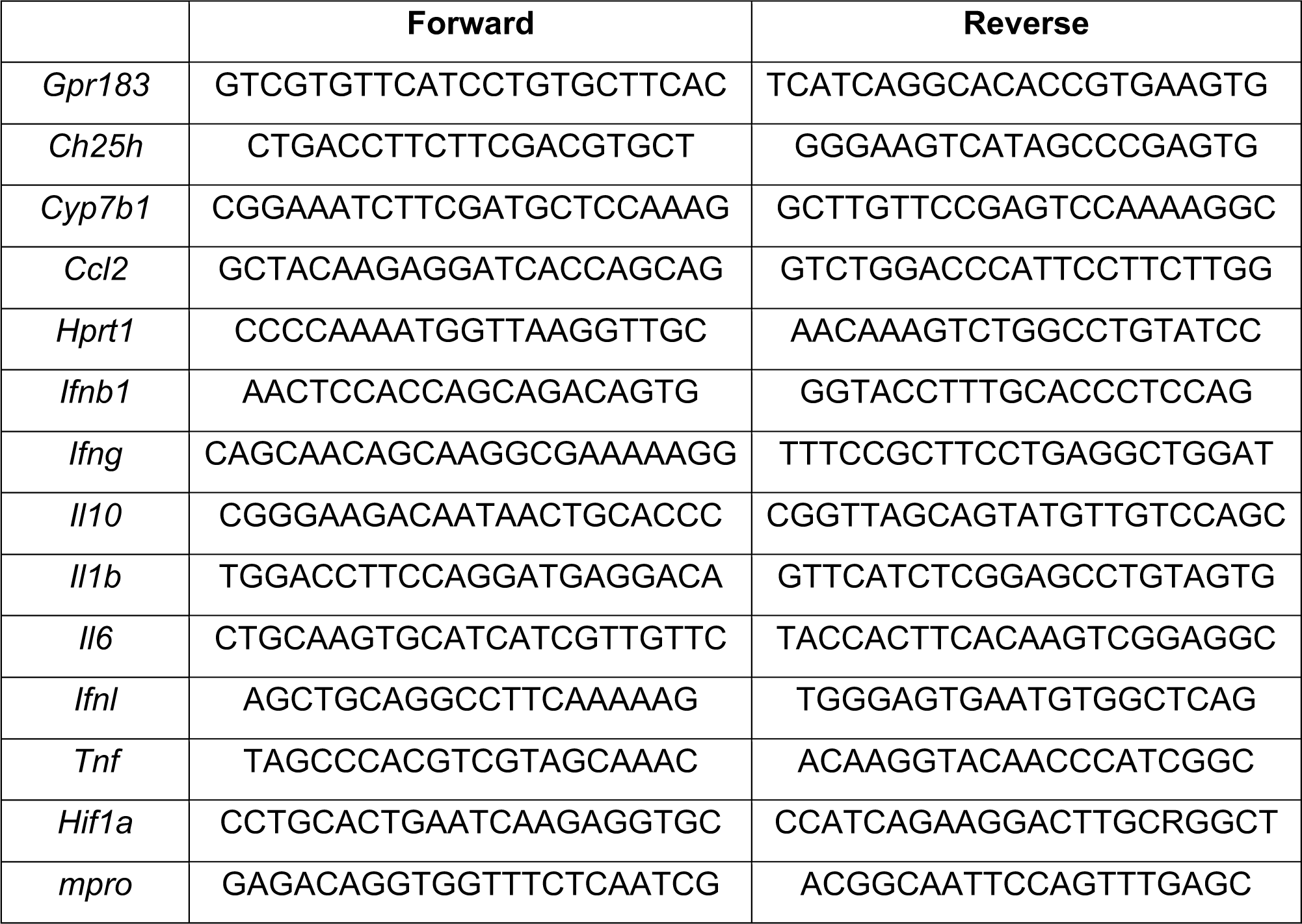
Primers used in this study.

